# Claudin-4 remodeling of nucleus-cell cycle crosstalk maintains ovarian tumor genome stability and drives resistance to genomic instability-inducing agents

**DOI:** 10.1101/2024.09.04.611120

**Authors:** Fabian R. Villagomez, Julie Lang, Daniel Nunez-Avellaneda, Kian Behbakht, Hannah L. Dimmick, Patricia Webb, Kenneth P. Nephew, Margaret Neville, Elizabeth R. Woodruff, Benjamin G. Bitler

**Affiliations:** Division of Reproductive Sciences, Department of Obstetrics and Gynecology, School of Medicine, University of Colorado, Anschutz Medical Campus, Aurora, Colorado; Department of Immunology and Microbiology, University of Colorado Anschutz Medical Campus, Aurora, USA; Deputy Directorate of Technological Development, Linkage, and Innovation, National Council of Humanities, Sciences, and Technologies, Mexico City, Mexico; Division of Gynecologic Oncology, Department of Obstetrics and Gynecology, The University of Colorado Anschutz Medical Campus, Aurora, Colorado; Medical Sciences, Indiana University School of Medicine, Bloomington, Indiana; Melvin and Bren Simon Comprehensive Cancer Center, Indiana University, Indianapolis, Indiana; Department of Anatomy, Cell Biology & Physiology, Indiana University, Indianapolis, Indiana

**Keywords:** Ovarian cancer, genomic integrity, nuclei, lamin B1, perinuclear F-actin, PARPi, forskolin, LAT1, HIF-1, ROS

## Abstract

During cancer development, the interplay between the nucleus and the cell cycle leads to a state of genomic instability, often accompanied by observable morphological aberrations. These aberrations can be controlled by tumor cells to evade cell death, either by preventing or eliminating genomic instability. In epithelial ovarian cancer (EOC), overexpression of the multifunctional protein claudin-4 is a key contributor to therapy resistance through mechanisms associated with genomic instability. However, the molecular mechanisms underlying claudin-4 overexpression in EOC remain poorly understood. Here, we altered claudin-4 expression and employed a unique claudin-4 targeting peptide (CMP) to manipulate the function of claudin-4. We found that claudin-4 facilitates genome maintenance by linking the nuclear envelope and cytoskeleton dynamics with cell cycle progression. Claudin-4 caused nuclei constriction by excluding lamin B1 and promoting perinuclear F-actin accumulation, associated with remodeling nuclear architecture, thus altering nuclear envelope dynamics. Consequently, cell cycle modifications due to claudin-4 overexpression resulted in fewer cells entering the S-phase and reduced genomic instability. Importantly, disrupting biological interactions of claudin-4 using CMP and forskolin altered oxidative stress cellular response and increased the efficacy of PARP inhibitor treatment. Our data indicate that claudin-4 protects tumor genome integrity by remodeling the crosstalk between the nuclei and the cell cycle, leading to resistance to genomic instability formation and the effects of genomic instability-inducing agents.

Graphical abstract
Claudin-4 plays a crucial role in remodeling the cytoskeleton, particularly influencing nuclear architecture. This remodeling appears to act as a regulatory mechanism, limiting the progression of ovarian cancer cells into the S-phase and correlating with reduced genomic instability in ovarian tumors. Moreover, since olaparib treatment triggered a cellular oxidative response, it is likely that this claudin-4-mediated remodeling contributes to resistance against the effects of the PARP inhibitor.

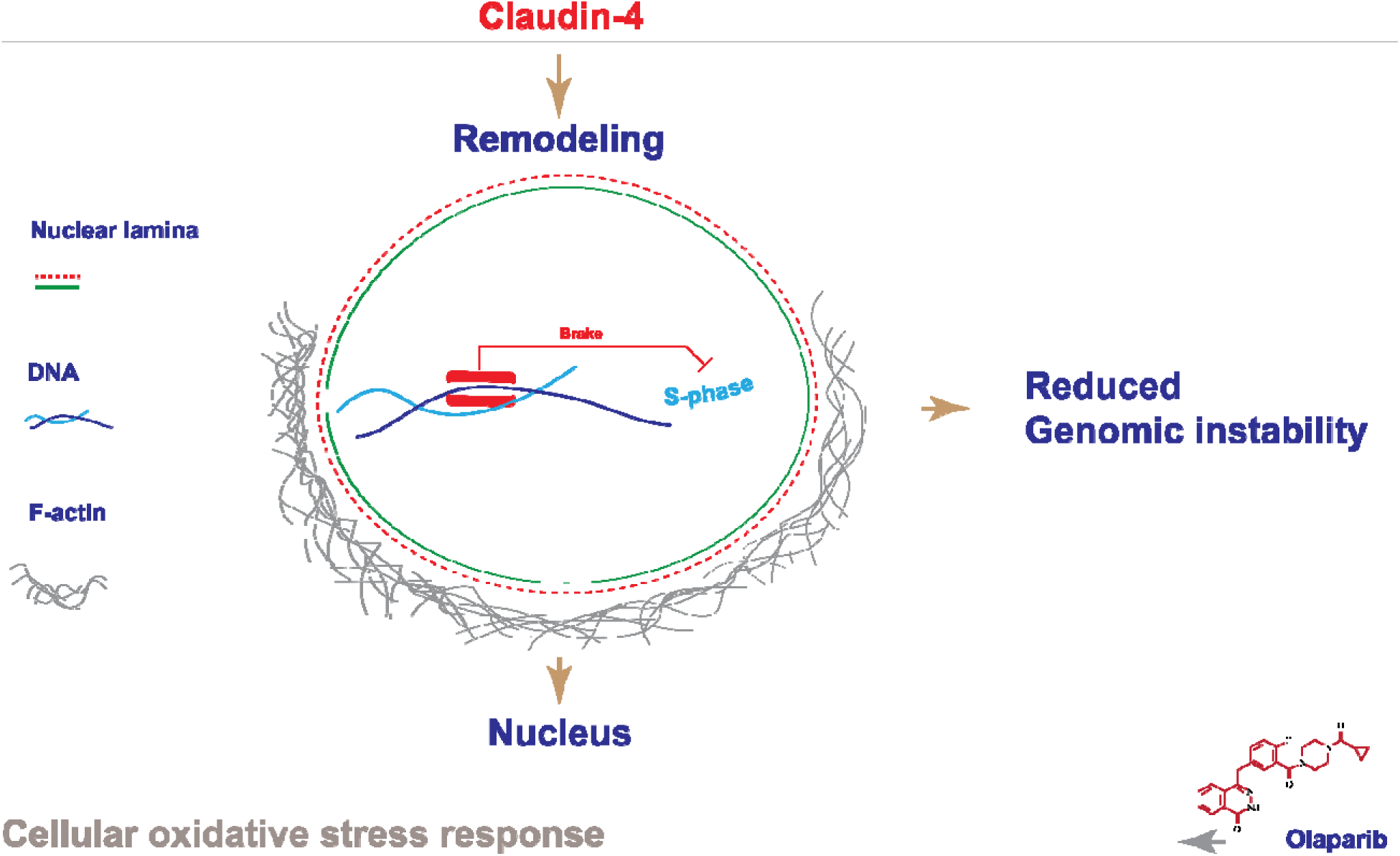

## INTRODUCTION

Cell cycle dysregulation is a fundamental hallmark of cancer that drives both altered cell proliferation and genomic instability, making it a critical therapeutic target.(1) The cell cycle plays a vital role in nuclear physiology, precisely coordinating nuclear envelope and cytoskeleton dynamics. This regulation ensures proper nuclear remodeling and safeguards genomic integrity throughout each phase of cell progression.(2–6) In addition, the physical connection of the nucleus with cell-to-cell junctions is a major component of the interplay between the cell cycle and nuclear physiology. This interplay occurs through the nuclear envelope, a membrane network composed of lamin proteins and surrounded by the LINC (Linkers of the nucleoskeleton to the cytoskeleton) complex, which binds to the cytoskeleton via actin, microtubules, and intermediate filaments.(3,6–11) Mechanistically, the connection between the nucleus and cell-to-cell junctions regulates both morphology and cell cycle progression through mechanotransduction, regulation of nuclear positioning and shape, spatial organization of tissues, and modulation of cell cycle checkpoints.(2,12–14) Moreover, significant alterations in nuclear morphology that are closely linked to genome instability have been observed during cancer development, (15,16) and tumor cells can maintain optimal tumor growth by either preventing or eliminating this hallmark of cancer (17–19)

Claudin-4 is aberrantly overexpressed in most epithelial ovarian carcinomas (EOC) (19–23) and is associated with resistance to therapy (23,24) and poor patient survival. This phenomenon is closely related to the regulation of genomic instability.(19,23) Claudin-4 is involved in many cellular functions, including cell proliferation (25) and DNA damage repair,(23) but it has been traditionally described as a cell-to-cell junction protein.(26,27). Recent research has shown that claudin-4 forms a functional axis with the amino acid transporters SLC1A5 and LAT1, playing a crucial role in controlling micronuclei, markers of genomic instability, through autophagy.(19) This indicates that claudin-4 actively participates in mitigating genomic instability after it arises.(19,23) Additionally, several studies underscore the potential clinical significance of claudin-4 in the treatment and prognosis of ovarian cancer.(19,20,23,26,28) However, the precise molecular mechanisms by which claudin-4 regulates genomic instability remain largely unexplored.

In this study we aimed to determine the influence of claudin-4 on cell cycle progression and nuclear architecture as well as how its regulation may lead to changes in genomic instability and therapy resistance in ovarian cancer cells. We targeted claudin-4 using a claudin mimic peptde (CMP) (DFYNP; 5 amino acids) (19,20,29). We aimed to disrupt interactions with claudin-4’s partner proteins and induce its mis-localization. (19,20,29,30) Additionally, we modulated claudin-4 expression in various EOC cells *in vitro*, including its overexpression in OVCAR8 cells and downregulation in OVCAR3 and OVCA429 cells, as reported previously.(19) We report a previously unknown “dual regulatory role” of claudin-4 in nuclear architecture and cell cycle progression, contributing to the crosstalk between nuclear physiology and the cell cycle. Furthermore, the dual role of claudin-4 was associated with ovarian tumor cell resistance to genome instability formation and to the effects of genomic instability-inducing agents, such as the PARP inhibitor olaparib.(31,32)

## METHODS

### Cell lines

Human derived cells, OVCA429 (RRID:CVCL_3936), OVCAR3 (RRID:CVCL_DH37), and OVCAR8 (RRID:CVCL_1629), collected from the Gynecologic Tissue and Fluid Bank, were cultured in RPMI-1640 medium (Gibco, Thermo Fisher Scientific, Cat. #11875) plus 10% heat-inactivated fetal bovine serum (Phoenix Scientific, Cat. # PS-100, Lot # 20055-01-01) and 1% penicillin/streptomycin (Corning, Cat. #30-002-CI) at 37°C and 5% CO2. HEK293-FT (RRID:CVCL_6911) were cultured similarly but in DMEM medium (Gibco, Thermo Fisher Scientific, Cat. #11995040).

### Vectors, lentivirus production and transduction

293FT cells were transfected using lipocomplexes (Lipofectamine 2000, ThermoFisher, cat: 11668-019) containing the viral packaging system of second generation (psPAX2, RRID:Addgene_12260; pMD2.G, RRID:Addgene_12259) as well as the lentiviral construct of interest (pLenti-Lifeact-tdTomato, RRID:Addgene_64048; pLenti-HIFR, RRID:Addgene_192946), respectively. Supernatant of transfected 293 FT cells was collected, filtered (0.45 µm), used, or stored (−80°C). Also, GFP-tubulin (EGFP-Tubulin-6, RRID:Addgene_56450) was cloned to the pCDH-CMV-MCS-EF1-Puro (EGFP-Tubulin-pCDH-CMV-MCS-EF1-Puro) vector using common cloning techniques (enzymatic restriction, NheI/BamHI; ligation, Plasmid sequencing) and viral particles were generated as indicated above.

### Cell cycle analysis by flow cytometry

2×10^5^ cells were seeded onto 6 well-plates (2mL RMPI complete medium). The next day, cells were washed (sterile PBS 1x) and media was changed (2mL RPMI complete medium). After 24h and 48h incubation, cells were washed (PBS 1X), detached (trypsin 0.25mM), and centrifuged (1500 rpm/5min). Cell pellets were resuspended in cold PBS 1X and centrifuged (1500 rpm/5min). Then, PBS was discarded, and cells were fixed using cold ethanol 70% (ethanol, milliQ water, v/v) for 30min at 4°C, then centrifuged (1500 rpm/5min/4°C). Afterwards, cells were washed twice with cold PBS 1X, and the PBS was discarded after centrifugation (1500 rpm/5min/4°C). Cells were treated with RNAse A (50µL/100µM) for 30min at room temperature (RT) and then stained with propidium iodide 300µL (50µM). Analysis was carried out in the Cancer Center Flow Cytometry Shared Resource (RRID: SCR_022035), University of Colorado Anschutz Medical Campus.

### Colony formation assay

3×10^4^ ovarian tumor cells were seeded onto 24 well-plates (1mL RPMI complete medium) and the next day, cells were washed with PBS 1X. Individual treatment (olaparib) or combination was applied (combination, combo: forskolin, fsk, 5µM; CMP, 400µM; olaparib, 600nM) in 1mL RPMI complete medium. Ovarian tumor cells were allowed to grow in the presence of individual or combination treatment for 7 days. OVCA429 cells received one dose of individual treatment due to more resistance to olaparib (at day 0; olaparib concentration: 120nM to 30µM), and OVCAR3/OVCAR8 cells received two doses of individual treatment due to less resistance to olaparib (at day 0 and day 4; olaparib concentration: 120nM to 1920nM). After 7 days of treatment cells were washed with PBS 1X and fixed (PSX 1X containing 10% acetic acid and 10% methanol) for 10 min and stained (using PBS 1X with 0.4% crystal violet and 20% ethanol for 10 min). To estimate the number of surviving cells, the cells were destained with PSX 1X containing 10% acetic acid and 10% methanol, and absorbance was read at 570 nm.

### Immunoblot

To analyze the protein expression levels of lamin B1, lamin A/C, LAT1, and Hif-1 alpha, tumor cells were scraped from culture plates in the presence of lysis buffer (30 mM Tris HCl pH7.4, 150 mM NaCl, 1% TritonX-100, 10% glycerol, 2 mM EDTA, 0.57 mM PMSF, 1X cOmplete™ Protease Inhibitor Cocktail), placed on a shaker for 10 minutes and spun at 13,000 rpm for 10 minutes. Protein was separated by SDS-PAGE and transferred to PVDF membrane using the TransBlot Turbo (BioRad). Membranes were blocked with Intercept Blocking Buffer (LI-COR, #927-60001) for 2 hours at RT. Rabbit anti-lamin B1 (Proteintech Cat# 12987-1-AP, RRID:AB_2136290, 1:3600), mouse anti-lamin A/C (Cell signaling Cat# 4777, RRID:AB_10545756, 1: 1000), rabbit anti-LAT1 (Cell signaling Cat# 5347, RRID:AB_10695104, 1: 1000), rabbit anti-hif-1 alpha (Proteintech Cat# 20960-1-AP, RRID:AB_10732601; 1: 1000), rabbit anti-GAPDH (Sigma Cat# HPA040067, RRID:AB_10965903, 1: 1000), mouse anti-β-actin (Abcam Cat# ab8226, RRID:AB_306371, 1: 5,000), primary antibody incubation was performed overnight at 4 °C. Membranes were washed 3 times for 5 minutes each in TBST (50 mM Tris pH 7.5, 150 RT temperature, followed by secondary antibodies for 2h at RT. Membranes were washed again 5 times for 5 min each in TBST. For fluorescent detection, bands were visualized using the LI-COR Odyssey Imaging System. CMP was synthesized as previously reported.(29)

### Immunofluorescence

Cells were fixed with paraformaldehyde at 4% (PBS 1X) for 10 minutes, followed by permeabilization (30 minutes, 0.1% Triton X-100, PBS 1X). Blocking was carried out by 2h incubation with BSA at 5% (PBS 1X, RT, shaking). Primary antibodies (as cited above, lamin B1, 1: 800; lamin A/C, 1: 100; LAT1, 1: 100; hif-1 alpha, 1: 100) were incubated (BSA at 2%, PBS 1X) overnight at 4°C/shaking. Secondary antibodies (AlexaFluor546 anti-mouse, ThermoFisher, cat: A-11030, at 2 µg/mL; AlexaFluor647 anti-rabbit, ThermoFisher, cat: A32733, at 2 µg/mL) were incubated 2h/shaking at RT (BSA at 2%, PBS 1X). Nuclei were stained with DAPI at 1 µg/mL (PBS 1X) for 10 minutes. All microscopy acquisition was carried out in the Neurotechnology Center, University of Colorado Anschutz Medical Campus.

### Live-cell imaging

2×10^5^ cells were seeded onto glass bottom dishes (35mm, No 1.5; MatTek, Cat. # P35G-1.5-14-C) and cultured in 2mL RMPI complete medium without phenol red (ThermoFisher, Cat. # 11835030). Nuclei were stained using Hoechst 1µM (ThermoFisher, 62249). All microscopy acquisition (FV1000, Olympus) was carried out in the Neurotechnology Center, University of Colorado Anschutz Medical Campus.

### Measurement of reactive oxygen species (ROS)

2×10^5^ cells were seeded onto 6 well-plates, and the following day, cells were washed with 1X sterile PBS and treated (combo: olaparib 600nM, forskolin, fsk, 5µM, and CMP 400 µM for 2h). TBHP was used as a positive control (250µM, 2h; all 2mL complete RPMI medium). To measure ROS, we used the DCFDA / H2DCFDA - Cellular ROS Assay Kit (Abcam, cat# ab113851). Cells were stained for ROS using DCFDA at 10µM (1mL RPMI medium final volume) for 40 minutes/37°C. Cells were put on ice and then analyzed via flow cytometry. All analyses were carried out in the Cancer Center Flow Cytometry Shared Resource, University of Colorado Anschutz Medical Campus.

### Statistical Considerations

ImageJ (NIH) and Prism software (v9.0) were used for microscopy and statistical data analysis, respectively. At least 3 independent experiments were conducted for most experiments. Unpaired t-tests and Mann–Whitney tests; Kruskal–Wallis test and one-way ANOVA with Dunn’s or Tukey’s multiple comparisons test were employed, based on normality data distributions and number of variables. The level of significance was p < 0.05.

### Data Availability Statement

All data is provided within the manuscript and supplemental information.

## RESULTS

### Claudin-4 enables ovarian tumor cells to modify the entry to and exit from cell cycle phases

Tumor heterogeneity is prevalent in cancer, even among tumors of the same type.(33) This heterogeneity is strongly linked to genomic instability and is associated with varying gene expression profiles and biological functions, particularly those related to the cell cycle. This variability is also evident in cell lines derived from tumors.(34,35) Given the association of claudin-4 with both the cell cycle(28) and genomic instability (19,23) in ovarian tumor cells, we evaluated cell cycle progression in different epithelial ovarian cancer cells (EOCs; OVCAR8, OVCA429, and OVCAR3). These cells were evaluated at 2 time points (24h and 48h post plating), followed by evaluation of the cell cycle. Assessment of these parental ovarian tumor cells indicated that all cell types were capable of transitioning through the cell cycle phases: G0-G1, S-phase, and G2-M. However, the cells tended to remain in the G0-G1 phase for extended periods, leading to diminished progression into the S and G2-M phases (*See Supplementary Figure 1a*) and leading to heterogeneous distributions of cells across the different cell cycle phases, with an increase in the proportion of cells in the G0-G1 phase over time (*See Supplementary Figure 1b*). This behavior may be linked to decreased nutrient availability over time, as suggested by previous studies.(19,36) Supporting this, we evaluated the cell cycle under reduced nutrient conditions in OVCAR8 cells, which have the fastest doubling time among the cancer cell lines studied. Our analysis revealed that ovarian tumor cells cultured under reduced nutrient conditions exhibited significantly reduced cell cycle progression (*See Supplementary Figure 1c*). Subsequently, we evaluated the effect of claudin-4 overexpression (in OVCAR8 cells, which do not express claudin-4) and downregulation (in OVCA429 and OVCAR3 cells, which do express claudin-4), as we previously reported,(19) on cell cycle progression. These manipulations led to modifications in the cell cycle progression when compared to wild-type (WT) cells. Claudin-4 overexpression was associated with a significant reduction in cells present in the S-phase (**Figure 1a**). At the same time, its downregulation resulted in significantly more cells present in the G2-M and fewer in the G0-G1 phases in OVCA429 cells (**Figure 1b**). Although it was not statistically significant, a similar pattern was observed in OVCAR3 cells at 24h (**Figure 1c**). A previous report also noted a higher number of cells in the G2-M phase during the downregulation of claudin-4 in OVCAR3 cells, which were synchronized by starvation at 6h.(28) Overall, our results show that claudin-4 significantly influences the progression of ovarian tumor cells through the cell cycle. Specifically, claudin-4 expression appears to arrest some ovarian tumor cells in the G0-G1 phase, resulting in fewer cells transitioning to the S-phase (**Figure 1a**). This finding is further supported by the observation that claudin-4 downregulation in OVCA429 cells leads to an increase in the number of cells in the G2-M phase and a decrease in the G0-G1 phase (**Figure 1b**). These findings suggest that claudin-4 is crucial for precisely controlling cell cycle phase transitions. Together, these responses suggest that claudin-4 overexpression enhances control over the cell cycle in ovarian cancer cells by slowing progression through the S-phase and ensuring proper entry and exit from the G2-M phase toward the G0-G1 phase (**Figure 1d**), potentially mitigating factors that could lead to genomic instability.

**Figure 1.**
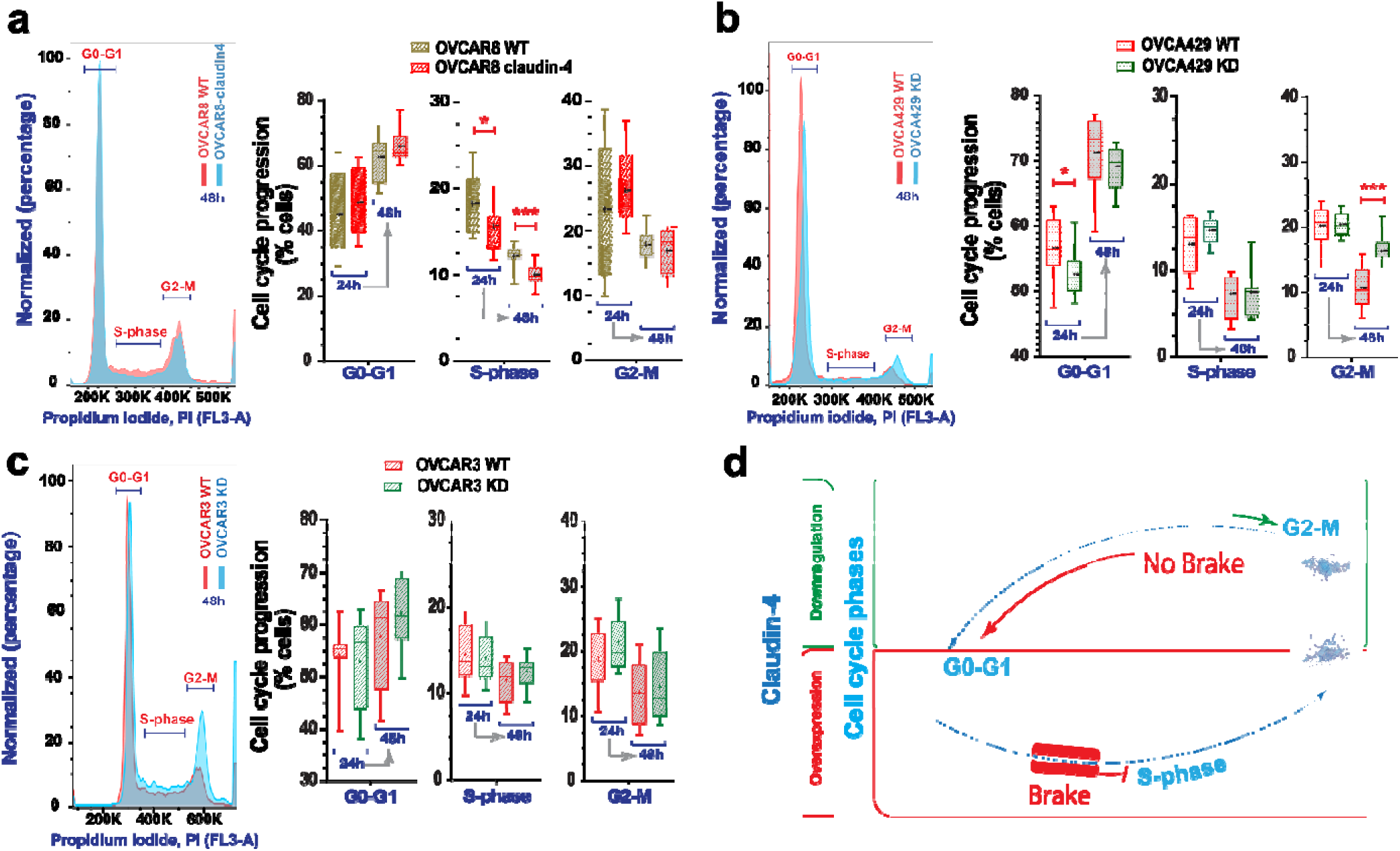
Cell cycle progression during claudin-4 modulation. Ovarian tumor cells were cultured and stained for propidium iodide (PI) at 24h and 48h, and cell cycle progression was evaluated via flow cytometry. (**a**) Representative histograms of cell cycle phases during claudin-4 overexpression; right, percentages of cells at each cell cycle stage. Effect of claudin-4 downregulation in OVCA429 **(b)** and OVCAR3 **(c)**. (**d**) Model illustrating that claudin-4 overexpression reduces the proportion of tumor cells in the S-phase of the cell cycle, while its downregulation results in an accumulation of cells in the G2-M phase and a decrease in the G0-G1 phase. (4 independent experiments; Two-tailed Unpaired t test; significance p<0.05). Graphs show mean and SEM (standard error of the mean).

### Claudin-4 remodels the nuclear architecture by influencing both the nuclear lamina and the actin-cytoskeleton dynamics

Genomic instability is intimately tied to nuclear dynamics during the cell cycle. Nuclear remodeling can give rise to significant morphological and cell cycle changes, which are common characteristics of cancer development and indicate genomic instability. (10,11,37,38) To further explore the role of claudin-4 in the cell cycle, we tested the effects of claudin mimic peptide (CMP) on ovarian cancer cells. The cells were then stained via immunofluorescence to mark the main components of the nuclear lamina (lamin B1 and lamin A/C) (39) and with phalloidin to label the actin-cytoskeleton, as previously reported.(40) We found significant remodeling of the nuclear architecture characterized by changes in the nuclear lamina and the actin-cytoskeleton. Specifically, claudin-4 overexpression led to reduced accumulation of lamin B1 in the nucleus (**Figure 2a**), while its downregulation significantly increased lamin B1 nuclear localization (**Figure 2b, c**). Notably, claudin-4 is known to be localized in the nucleus of ovarian tumor cells,(26) where we observed an inverse relationship with lamin B1 (**Figure 2a-c**). Given that no significant changes were detected in the expression levels of lamin B1 or lamin A/C (*See Supplementary Figure 1b-d*), it appears that claudin-4 primarily affects the localization of lamin B1.(39) Targeting claudin-4 via CMP treatment affected the nuclear lamina as well, causing significant changes in the nuclear localization of both lamin B1 and lamin A/C (**Figure 2a-c**). In OVCAR8 WT cells, treatment with CMP led to an increased nuclear localization of lamin A/C but not lamin B1. However, this CMP effect on Lamin A/C was reversed in claudin-4 overexpressing cells (**Figure 2a**), indicating a specific role of claudin-4 in modulating the CMP- induced changes in lamin A/C distribution. Additionally, in OVCA429 WT cells, which naturally express claudin-4, CMP treatment was associated with a decreased accumulation of lamin B1. In contrast, we observed an increase in both lamin B1 and lamin A/C levels in OVCAR3 WT cells. Thus, the effect of CMP on nuclear lamina components (lamin B1 and lamin A/C) varied among different ovarian cancer cell lines. This variation suggests that the proteins interacting with claudin-4 in these cells may differ, potentially influencing CMP’s ability to target claudin-4’s role in the nuclear lamina. (19,26,29) Our data indicate that claudin-4 expression and function play a dominant role in the modification of nuclear lamina components. (**Figure 2**)

**Figure 2.**
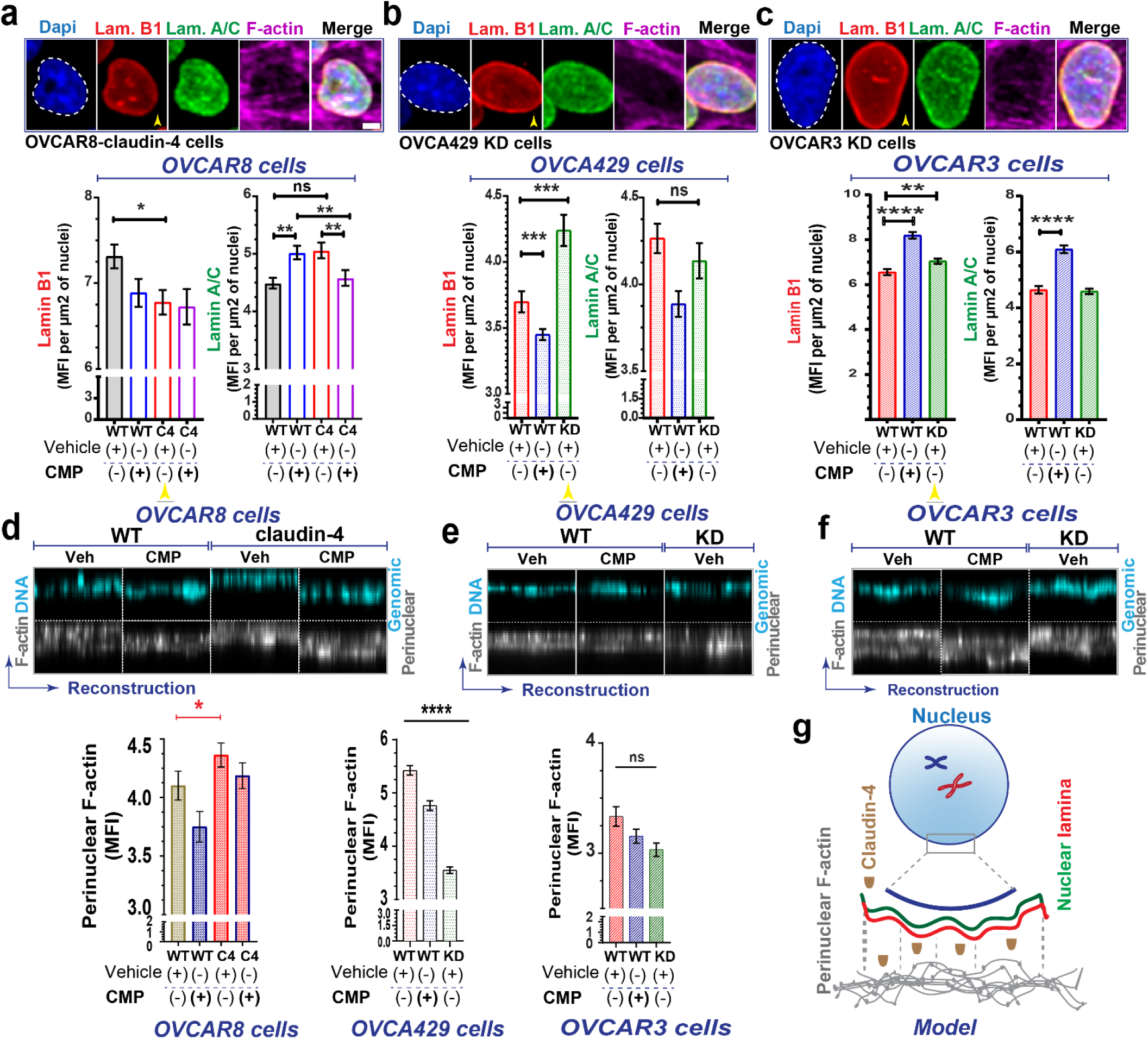
Claudin-4 modifies the nuclear architecture of ovarian tumor cells. The association of claudin-4 with the actin cytoskeleton was evaluated in fixed and living cells. Ovarian tumor cells were treated with CMP (400µM) for 48 h and were stained via immunofluorescence to mark lamin B1 and lamin A/C, while the actin-cytoskeleton was stained with phalloidin. We then carried out a morphometric characterization. **(a)**. Top, selected confocal images (maximum projections) corresponding to OVCAR8 cells overexpressing claudin-4 and claudin-4 downregulation in OVCA429 cells (bottom) (**b**). Results of similar experiments in OVCAR3 claudin-4 KD cells (**c**), showing the nuclear lamina components (lamin B1, and lamin A/C), bottom, with corresponding quantification of nuclear accumulation of the nuclear lamina components (yellow arrowheads highlight comparison of lamin B1 during claudin-4 overexpression and downregulation) under different conditions (three independent experiments; Kruskal-Wallis test with Dunn’s multiple comparisons, p<0.05). **(d)**, **(e)**, and **(f)**, top, show reconstructions of perinuclear F-actin and genomic DNA (from confocal z-stacks), and, bottom, corresponding quantification in OVCAR8, OVCA429, and OVCAR3 cells, respectively. Additionally, a drawing highlights the remodeling effect of claudin-4 in the nuclear architecture (n= OVCAR8, 1711 cells; OVCA429, 2630 cells; OVCAR3, 2365 cells; Two-tailed Mann Whitney test, Kruskal-Wallis test with Dunn’s multiple comparisons). Graphs show mean and SEM, scale bar 5µm.

Polymeric actin (F-actin) was also affected during claudin-4 modulation, particularly in the perinuclear region and the cytoplasmic fibers (*See Supplementary Figure 2*). F-actin was more localized in the perinuclear region during claudin-4 overexpression (**Figure 2d**) and exhibited the opposite effect during its downregulation, especially in OVCA429 cells (**Figure 2e, f**). Additionally, all ovarian tumor cells treated with CMP demonstrated a pattern of reduced perinuclear F-actin localization, implying that both expression and proper claudin-4 localization are required perinuclear F-actin positioning. We observed a positive correlation between claudin-4 expression and perinuclear F-actin accumulation (**Figure 2d-f**). This, along with the observed inverse relationship between claudin-4 and lamin B1 (**Figure 2a-c**), suggest a competitive exclusion mechanism,(41,42) where claudin-4 regulates lamin B1 dynamics. Overall, our results indicate that claudin-4 remodels nuclear architecture by changing the dynamics of lamin B1 and perinuclear F-actin (**Figure 2g**), which could provide an advantage for tumor cells. In addition, when claudin-4 was downregulated, we detected significantly lower concentrations of F-actin at the cell-to-cell junctions (junctional actin) (**Figure 3, a-b**). However, CMP treatment did not affect this accumulation, indicating that maintenance of junctional F-actin is independent of claudin-4 inhibition compared to perinuclear F-actin. We hypothesized that claudin-4’s regulation of perinuclear F-actin is more dominant than its regulation of junctional F-actin. To explore this hypothesis, we generated ovarian tumor cells expressing LifeAct (a marker of F-actin). As previously reported, we performed time-lapse confocal imaging and kymographs to capture the temporal motion of junctional F-actin.(40,42) Upon downregulation of claudin-4, the cellular connections between ovarian tumor cells exhibited increased mobility and progressively became more irregular. This shift underscores significant alterations in the plasticity of cell-to-cell interactions. (**Figure 3c, d***, Supplementary video 1 and 2*). This observation aligns with results detected in fixed cells (**Figure 3a, b**) and strongly suggests that claudin-4 participates in the plasticity of cell-to-cell interactions, potentially regulating the connections between tumor cells to promote survival in malignant ascites.(42,43)

**Figure 3.**
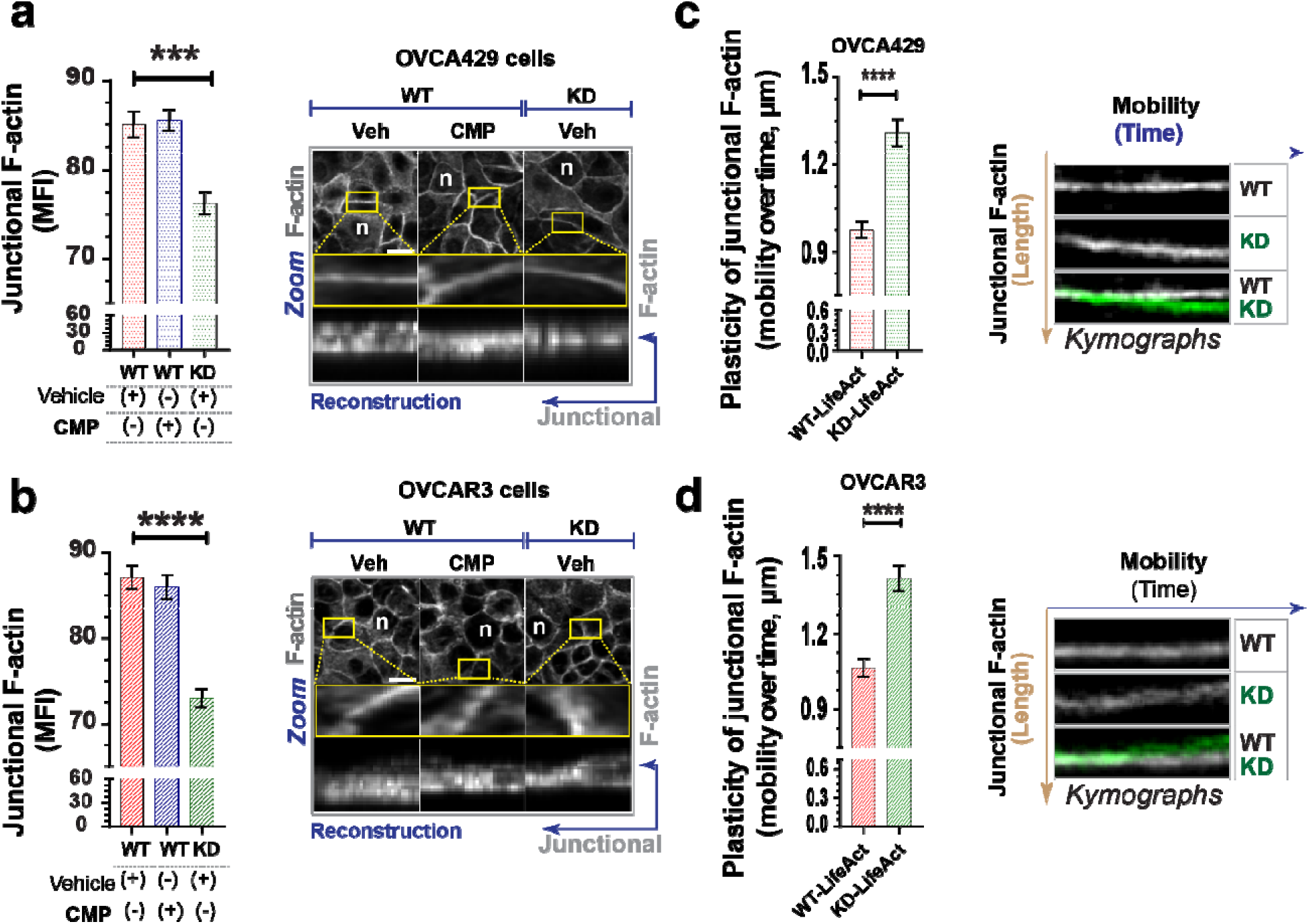
Disruption of claudin-4 impacts dynamics of junctional actin. Ovarian tumor cells were treated with CMP (400µM) for 48h and stained to mark the actin-cytoskeleton. Additionally, ovarian tumor cells expressing LifeAct (a marker of F-actin) were used to visualize actin dynamics in living cells. **(a)** and **(b)**, left, quantification of junctional F-actin from reconstructions (from confocal z-stacks); right, confocal images (maximum projection) and zoom, followed by reconstruction of selected regions of interest, ROIs (at junctional F-actin) from OVCA429 cells and OVCAR3 cells (bottom), respectively (n= OVCA429, 783 cells; OVCAR3, 825 cells; Kruskal-Wallis test with Dunn’s multiple comparisons). (**c**) and **(d),** kymographs illustrating the movement of junctional F-actin (vertical brown arrow) over time (horizontal blue arrow), generated from different regions of interest (ROIs) during confocal live-cell imaging of transduced OVCA429 cells (n=142) with LifeAct to mark F-actin (without any stimuli and cultured for 24h) and OVCAR3 cells (bottom) (n=116), respectively (Two-tailed Mann Whitney test) (3 independent experiments; significance, p<0.05). Graphs show mean and SEM.

Cell-to-cell junctions are known to be physically connected with the nuclear lamina and perinuclear F-actin through the cytoskeleton,(6,44) which influences the positioning of the nucleus for proper cell cycle progression.(6,44,45) Given that CMP treatment affected the nuclear lamina and perinuclear F-actin, we hypothesized that the mobility of the nucleus could also be affected by claudin-4 manipulation. We evaluated nuclear mobility during CMP treatment in living OVCA429 WT cells and observed that CMP (sequence: D<u>F</u>YNP) significantly reduced nuclear mobility compared to a control peptide (sequence: D<u>G</u>YNP) (*See Supplementary Figure 3a*). This effect underscores the importance of the conserved sequence targeted by CMP (20,29) for disrupting claudin-4’s association with nuclear structures like the nuclear lamina and perinuclear F-actin. The reduced nuclear mobility might result from decreased cytoskeletal connections to the nucleus. (3,6,44) Supporting this concept, confocal live cell imaging of ovarian tumor cells expressing LifeAct and GFP-tubulin revealed that claudin-4 downregulation in OVCA429 cells altered F-actin levels and microtubule network formation (*See Supplementary Figure 3b, c*). This finding is consistent with earlier reports showing that claudin-4 influences tubulin polymerization,(28) reinforcing its close association with the cytoskeleton in ovarian tumor cells. Claudin-4 may influence the polymerization of actin and tubulin, which is crucial for regulating nuclear connectivity (*See Supplementary Figure 3d*). The data highlight a crucial claudin-4-mediated interplay between the cell cycle and nuclear architecture, possibly impacting the mitotic capabilities of ovarian tumor cells.(46,47) Specifically, ovarian tumor cells that overexpress claudin-4 may experience delays in nuclear remodeling, prolonging the transition to the S-phase of the cell cycle, and resulting in fewer cells progressing to the S-phase, potentially reducing the likelihood of genetic instability.(48,49)

### Overexpression of claudin-4 leads to constriction in nuclear size, which is associated with a reduction in genomic instability

To further explore the association of claudin-4 with genomic instability, we analyzed human ovarian tumors in The Cancer Genome Atlas. We correlated levels of claudin-4 expression with chromosomal copy number alterations as an indicator of genomic instability. Ovarian tumors with high claudin-4 expression showed greater than 2-fold reduction in genomic instability compared to tumors with low claudin-4 expression (2.32% [95% CI 2.28-2.36] vs. 5.00% [95% CI 4.93-5.08], P<0.0001) (**Figure 4a**), consistent with previous reports indicating an association between claudin-4 expression and decreased genetic mutations.(23) Also, claudin-4 has been previously identified as a key regulator of micronuclei through autophagy,(19) further confirming its role as a regulator of genetic instability.

**Figure 4.**
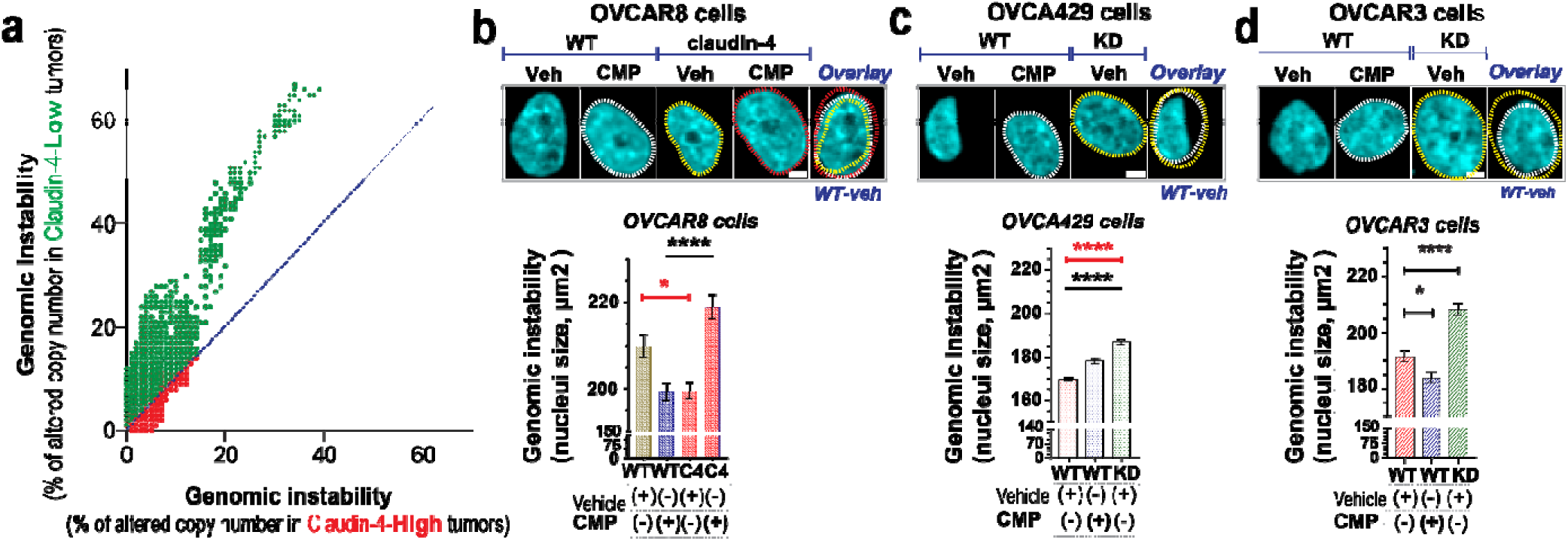
Claudin-4’s association genomic instability correlates with nuclei constriction. The association of claudin-4 with genomic instability was analyzed in human tumors (TCGA) and *in vitro* by treating cells with CMP (400µM) for 48h followed by single cell analysis of fixed cells (stained with DAPI to mark DNA). **(a)**, correlation of genomic instability (indicated as % of altered chromosomic copy numbers) in human ovarian tumors associated with levels of claudin-4 expression. **(b)**, top, confocal images showing (maximum projections) genomic instability (indicated as nuclei size, bottom: corresponding quantification) associated with claudin-4 overexpression or downregulation (knockdown, KD) in OVCA429 **(c)** and OVCAR3 **(d)**. (n= OVCAR8, 1711 cells; OVCA429, 2630 cells; Two-tailed Mann Whitney test, Kruskal-Wallis test with Dunn’s multiple comparisons). (3 independent experiments; significance, p<0.05. Graphs show mean and SEM, scale bar 5µm.

Nuclear size and chromosomal amplifications are positively correlated in ovarian cancer cells.(15,16,50–53) Therefore, we investigated the connection between claudin-4 expression and nuclear size by morphological characterization of nuclei from ovarian tumor cells stained with DAPI during claudin-4 disruption. We found that the nuclear size was decreased in claudin-4 overexpressing cells, and this phenotype was reversed by CMP treatment (**Figure 4b**). Conversely, nuclear size expanded when claudin-4 was downregulated (**Figure 4c, d**). However, CMP only increased nuclear size in OVCA429 cells, while in OVCAR3 cells, the reverse effect occurred. Although it is clear that claudin-4 plays a role in regulating nuclear size and that CMP can moderate its effects, this finding highlights the intertumoral diversity of ovarian tumor cells.(19,54) It is possible that claudin-4-interacting proteins in the plasma membranes of OVCA429 and OVCAR3 differ, leading to differences in CMP targeting effects. (19) Additionally, the varying levels of perinuclear F-actin observed in OVCA429 and OVCAR3 cells during CMP treatment further highlight these differences (**Figure 2e, f**). The nucleus is typically larger in G2-M than in S or G1-G0 phases.(55) We observed a correlation between claudin-4 modulation’s impact on nuclear size and its influence on cell cycle progression. Claudin-4 overexpression and its downregulation correlated with nuclei constriction and expansion, respectively (**Figure 4b-d**), which aligns with a reduced number of cells observed in the S-phase during claudin-4 overexpression (**Figure 1a**). Conversely, more cells were noted in the G2-M phase during claudin-4 downregulation (**Figure 1b**). Thus, in our *in vitro* models, more claudin-4 expression leads to a slowed progression through the cell cycle, possibly allowing repair of DNA damage or increased regulation of chromosomal separation (**Figure 4a**).(23) These findings potentially underscore the clinically significant effect of claudin-4 in ovarian tumors (23) through control of the cell cycle and maintenance of genome integrity.

### Enhancing the efficacy of a PARP inhibitor by disrupting claudin-4-functional effects in ovarian tumor cells

Our results suggest that claudin-4 protects ovarian tumors by modifying the interplay between the nuclear structure and the cell cycle through its close association with the cytoskeleton, ultimately reducing genomic instability (**Figure 4a**). High expression of claudin-4 predicts poor patient survival related to the development of therapy resistance (*See Supplementary Figure 4a*). To better understand the role of claudin-4 in therapy resistance, we evaluated the effect of claudin-4 modulation (overexpression and downregulation) during PARP inhibitor (olaparib) treatment. This treatment is known for inducing catastrophic genomic instability in BRCA-deficient tumors such as ovarian and breast cancer, (31,32) but it induces tumor cell death regardless of such mutations.(56–60) We reasoned that the reduction of genomic instability and slowing of mitosis by claudin-4 may interfere with olaparib effects on ovarian tumor cells. We evaluated the resistance of different ovarian tumor cell lines (OVCAR8 cells, BRCA1/2 no alteration; OVCA429 cells, BRCA1/2 no available data; OVCAR3 cells, BRCA1 no alteration, BRCA2 deep deletion)(19,54) by measuring colony formation over 7 days in the presence of varying doses of olaparib, finding that the cancer cells showed varying degrees of resistance to olaparib. OVCAR3 cells were the most sensitive to the higher concentrations used (15000nM to 30000nM), followed by OVCAR8 (*See Supplementary Figure 4b, c*), and then OVCA429 cells (**Figure 5**). Due to high sensitivity of OVCAR8 and especially OVCAR3 to olaparib, we used lower concentrations of olaparib for these cells than OVCA429 cells to evaluate the effect of targeting claudin-4 on resistance to this PARP inhibitor. The overexpression of claudin-4 in OVCAR8 cells (ovarian cancer subtype: low-grade serous ovarian carcinoma, LGSOC)(54) did not increase resistance to olaparib (**Figure 5a**). Similarly, claudin-4 downregulation in OVCA429 cells (ovarian cancer subtype: mucinous ovarian carcinoma, MOC)(54) was not linked to reduced resistance to olaparib (**Figure 5b**). Although the observation that OVCA429 cells treated with olaparib suggests increased resistance in claudin-4 KD cells compared to WT cells, it is also linked to a significant increase of OVCA429 claudin-4 KD cells in the G2-M phase of the cell cycle. (**Figure 1d**). Conversely, claudin-4 downregulation in OVCAR3 cells (ovarian cancer subtype: high-grade serous ovarian carcinoma, HGSOC) significantly decreased resistance to olaparib at lower doses (120nM and 240nM) (**Figure 5c**), consistent with prior findings.(23) This suggests that targeting claudin-4 could reduce the minimum concentration of olaparib required to inhibit HGSOC tumor growth.

**Figure 5.**
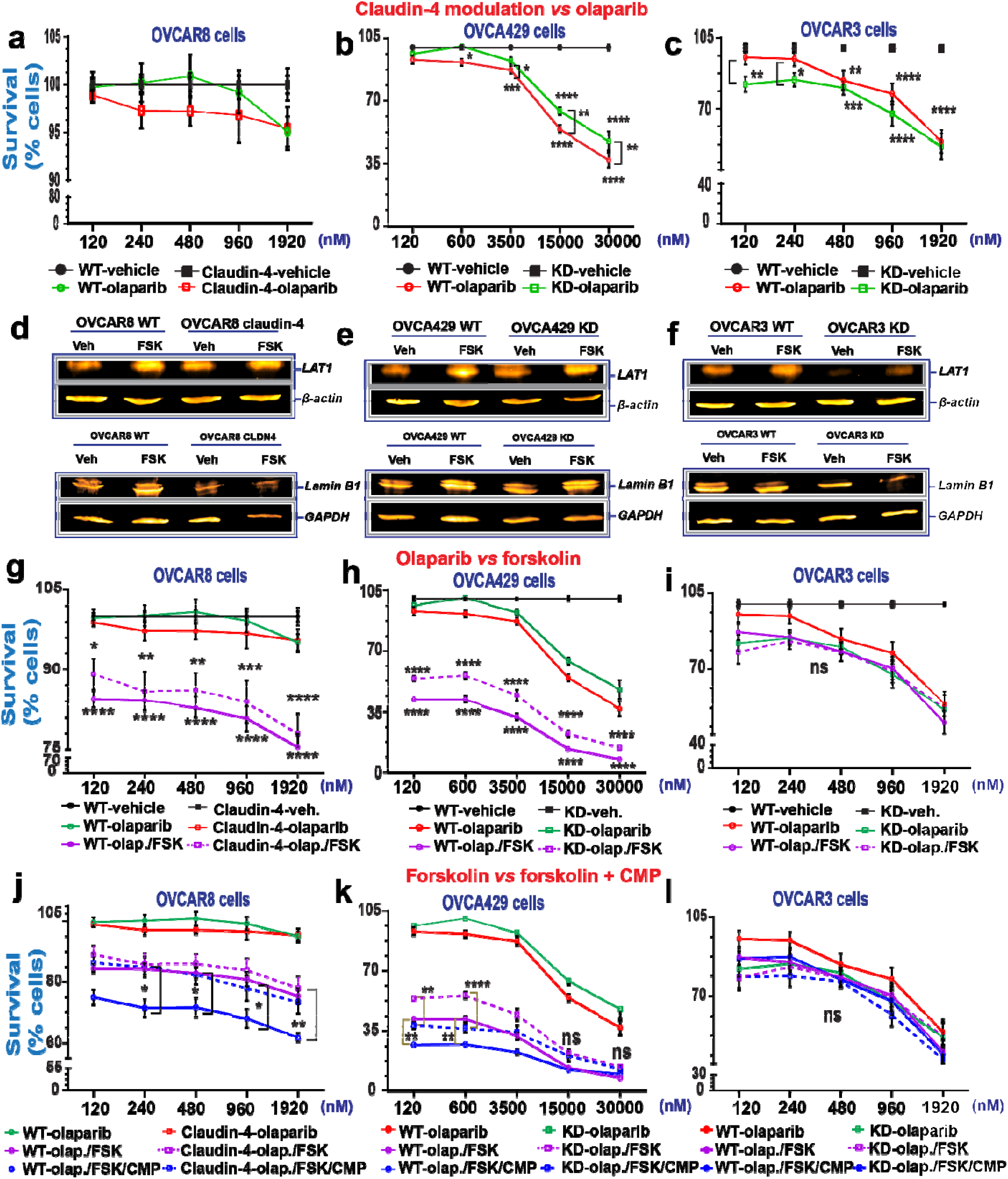
Targeting ovarian tumor cells during claudin-4 expression via forskolin and CMP to overcome resistance against olaparib. Survival of ovarian tumor cells was analyzed using the 7 day colony formation assay and two cycles of treatment (at day 0 and day 3) for OVCAR8 and OVCAR3 cells, and one treatment for OVCA429 (at day 0). (**a**), percentage of tumor cell survival during olaparib treatment and claudin-4 overexpression, and similar information during claudin-4 downregulation in OVCA429 cells (**b**) and OVCAR3 cells (**c**). Immunoblotting for LAT1 and lamin B1 during forskolin treatment (5µM/ 48h) during claudin-4 overexpression in OVCAR8 cells (**d, and bottom**), and similar information during claudin-4 downregulation in OVCA429 cells (**e, and bottom**) and OVCAR3 cells (**f, and bottom**). (**g**), percentage of tumor cell survival during olaparib treatment *vs* olaparib + FSK (5µM) and claudin-4 overexpression, and similar information during claudin-4 downregulation in OVCA429 cells (**h**) and OVCAR3 cells (**i**). (**j**), percentage of tumor cell survival during olaparib + FSK (5µM) *vs* olaparib + FSK (5µM) + CMP (400µM) and claudin-4 overexpression, and similar information during claudin-4 downregulation in OVCA429 cells (**k**) and OVCAR3 cells (**l**). (3 independent experiments; Two-way ANOVA; significance p<0.05). Graphs show mean and SEM.

Furthermore, we recently reported that claudin-4 forms a functional link with LAT1, an amino acid transporter, to regulate a form of genomic instability, micronuclei, via autophagy.(19) Thus, to leverage both claudin-4’s regulation of the actin cytoskeleton and LAT1 activity, we next evaluated with the use of forskolin (FSK), a compound that both modifies actin polymerization(61–63) and increases the expression of LAT1.(64) FSK also functions to increase levels of cAMP through the activation of adenyl cyclase.(65) Critically, FSK has been reported to have anti-ovarian tumor activity by improving the efficacy of a PARP inhibitor.(60) Consequently, we evaluated the effect of this compound on claudin-4’s functional influence in ovarian cancer cells. First, we confirmed the anti-tumor activity of FSK in OVCA429 WT cells (*See Supplementary Figure 4d*). Subsequently, we measured the protein expression level of LAT1 in ovarian cancer cells treated with FSK and confirmed that FSK upregulates the expression of LAT1(**Figure 5d-f**). This suggests that the reported link between claudin-4 and LAT1 could be altered due to the effect of FSK.(19) Also, we observed increased expression of lamin B1 during the same treatment in OVCAR8 WT and OVCA429 WT cells (**Figure 5d-f, bottom**). Thus, FSK treatment altered the expression of proteins functionally associated with claudin-4, such as LAT1(19) and lamin B1 (**Figure 2**). Next, we performed the same experiment as before, with the addition of a combined olaparib/FSK treatment. This combination treatment resulted in a more significant decrease in ovarian cancer cell survival than olaparib only, especially in OVCAR8 and OVCA429 cells (**Figure 5g-i**), suggesting that FSK enhances the efficacy of olaparib treatment. However, this enhancement of olaparib efficacy was not observed in OVCAR3 cells (**Figure 5i**). The muted effect of the combined treatment on OVCAR3 cells may be due to the amplification of KRAS, unlike the other ovarian cancer cells.(54) Intracellular levels of cAMP are increased by FSK treatment,(65) which interact with MAPK signaling.(66) KRAS is a master regulator of MAPK signaling,(67) and can lead to sustained MAPK signaling,(68,69) which may oppose the effects of FSK, causing the treatment to be ineffective in OVCAR3 cells.

In OVCAR8 claudin-4 overexpressing cells, the efficacy of olaparib/FSK was reduced compared to WT cells (**Figure 5g**), suggesting that claudin-4 overexpression may increase resistance to olaparib/FSK therapy. Therefore, we performed an experiment that included a triple combination of olaparib, FSK, and CMP in the colony formation assay. We confirmed that the combination of FSK and CMP maintain anti-proliferation activity before applying the combination treatment to all ovarian cancer cells (*See Supplementary Figure 4e*). Importantly, all ovarian cancer cells treated with the triple combination therapy exhibited significantly reduced survival at the lowest concentration of olaparib used (120nM) compared to the vehicle control. Notably, the combination therapy of olaparib, FSK, and CMP led to a substantial decrease in cancer cell survival in OVCAR8 WT and OVCA429 WT and KD cells compared to the olaparib/FSK treatment alone (**Figure j, k**). Conversely, in OVCAR3 cells, the tripartite combination treatment did not further decrease cancer cell survival compared to olaparib/FSK or olaparib alone (**Figure 5l**). These results highlight how claudin-4’s influence on nuclear architecture and the cell cycle contributes to resistance to olaparib. Thus, claudin-4 could represent a meaningful therapeutic target to decrease olaparib resistance and promote ovarian cancer cell death.

To gain further insight into the molecular mechanisms by which disrupting claudin-4 functionality leads to increased ovarian cancer cell death in response to olaparib, potentially involving alterations in LAT1 and lamin B1, we assessed the expression of these proteins at various time points during claudin-4 modulation (both overexpression and downregulation) and triple treatment (olaparib/FSK/CMP). We observed consistent increases in the amino acid transporter LAT1 expression across all ovarian cancer cells tested over time. Notably, in response to the triple treatment, LAT1 expression showed increases during claudin-4 overexpression in OVCAR8 cells, while it exhibited the most pronounced decreases during claudin-4 downregulation in OVCA429 and OVCAR3 cells (**Figure 6a-c**). These findings suggest an association between claudin-4 and LAT1 in olaparib resistance in ovarian cancer cells. Although lamin B1 expression was not consistent over time and showed more variations (*See Supplementary Figure 5a-c*), its intracellular distribution exhibited clear differences due to our tripartite treatment. For example, this treatment led to the formation of nuclear lamina blebs in OVCAR8 and OVCAR3 cells, while in OVCA429 cells, it correlated with a reduced accumulation of cytoplasmic puncta, both of which were enriched with lamin B1 (*See Supplementary Figure 5d-i*). Thus, it appears that the combination treatment (olaparib/FSK/CMP) disrupts LAT1 expression and the intracellular distribution of lamin B1, which correlates with decreased survival of ovarian cancer cells.

**Figure 6.**
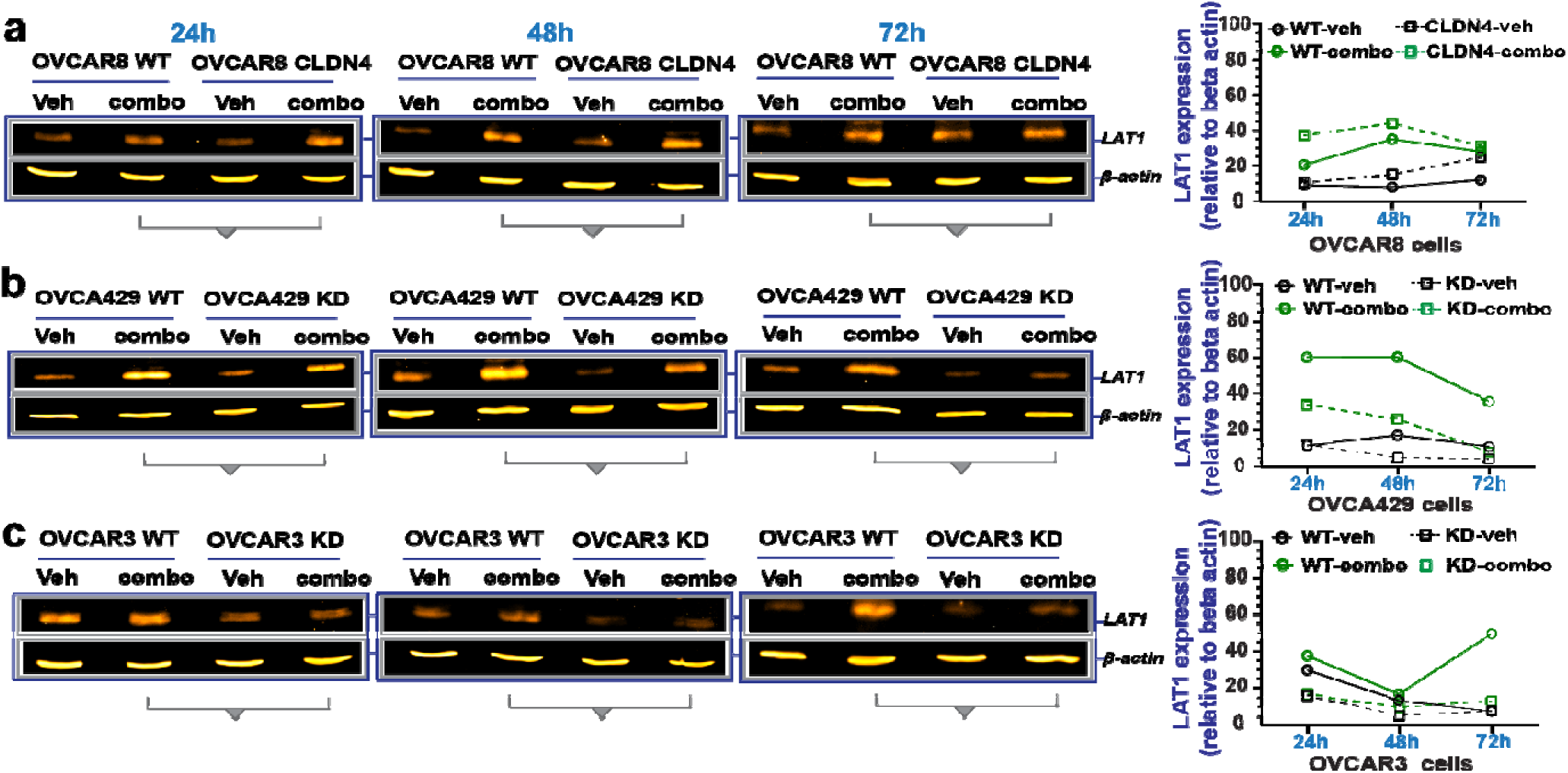
Effect of the combination treatment with olaparib, forskolin, and CMP on LAT1 expression. Ovarian tumor cells were treated with a tripartite combination of olaparib (600nM), fsk (5µM), and CMP (400µM) for different time points. Subsequently, cell lysates were obtained to carry out immunoblotting for LAT1. (**a**), (**b**), and (**c**) show LAT1 protein expression at different time points in OVCAR8, OVCA429, and OVCAR3 cells, respectively. On the right are graphs showing corresponding quantification of LAT1 from (**a), (b),** and **(c),** relative to loding control.

### Perturbing oxidative stress response has the potential to further reduce claudin-4-mediated olaparib resistance in ovarian tumor cell

In the cells treated with the triple combination, we observed a significant increase in reactive oxygen species (ROS) generation and hypoxia-inducible factor-1 α (hif-1 alpha) expression (**Figure 7a, b**) (*See Supplementary Figure 6a-c*). These findings indicate an upregulation of hypoxia-related elements,(70,71) suggesting an oxidative stress response to counteract the treatment effects.(72,73) Hypoxia, common in tumors and prolonged cell cultures,(32,74,75) allows tumor cells to adapt to low-oxygen environments,(72,73) contributing to therapy resistance.(76,77) This condition triggers hif-1 alpha, a key oxygen sensor,(70) and ROS production, (71) leading to an oxidative stress response.(78,79) Hif-1 alpha and ROS are interrelated in this stress response,(80) and tumor cells induce ROS production in response to PARP inhibitors.(60,81) Interestingly, proteins associated with claudin-4’s functional effects, such as lamin B1 (**Figure 2**) and LAT1(19) participate in cellular oxidative stress responses.(82–84).

**Figure 7.**
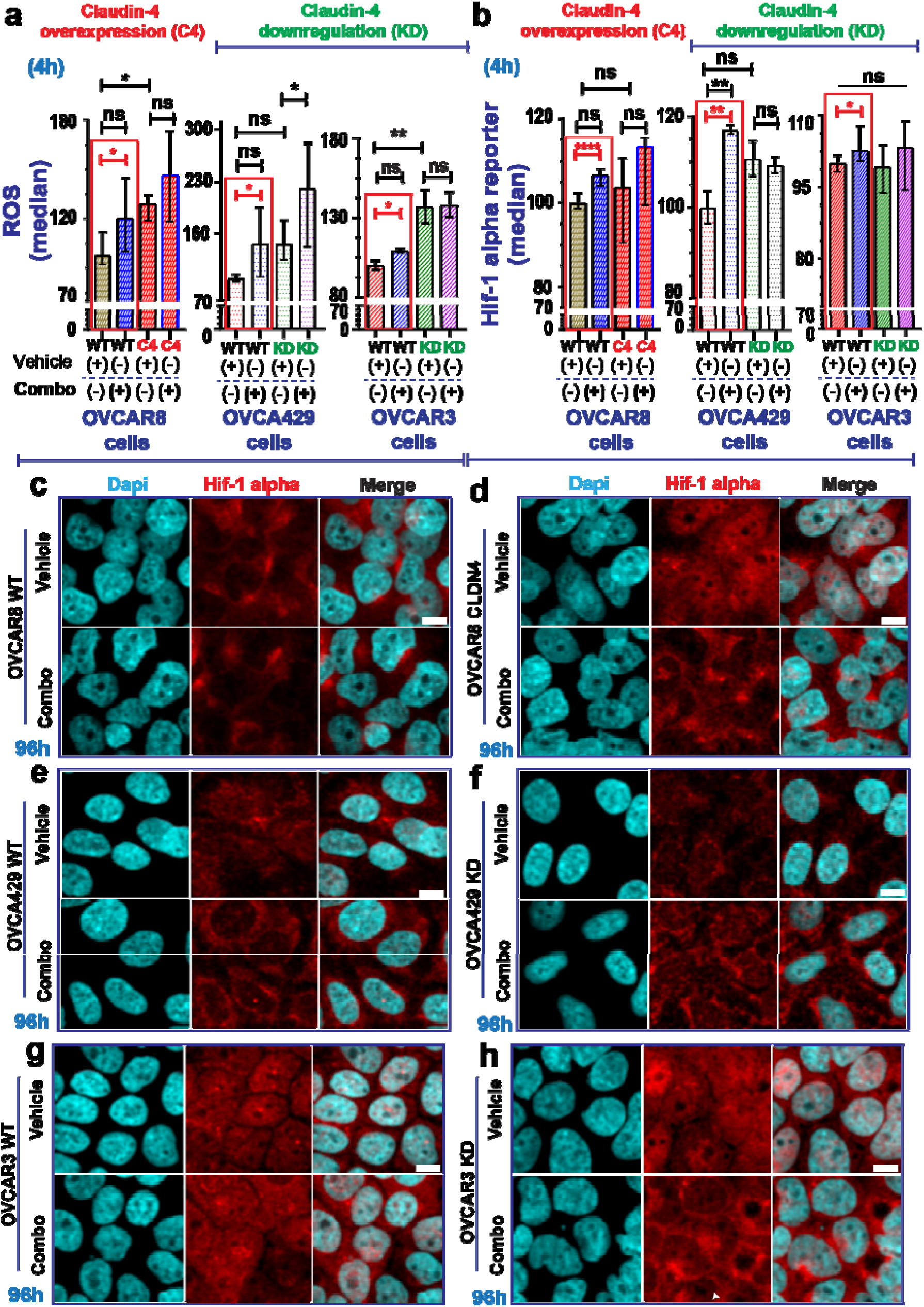
Cellular stress response evaluation in ovarian tumor cells treated with olaparib, forskolin, and CMP. Ovarian tumor cells were treated with a tripartite combination of olaparib (600nM), fsk (5µM), and CMP (400µM) for 4h to analyze reactive oxygen species (ROS) as well as a reporter gene for hif-1 alpha via flow cytometry. The same cells were treated similarly for 96h and then stained using immunofluorescence to mark lamin B1. ROS generation is indicated as normalization relative median of OVCAR8 WT, OVCA429 WT, and OVCAR3 WT cells without treatment (**a**) (2 independent experiments; Unpair t-test, red rectangle; One-way ANOVA and Tukey’s multiple comparison test, p<0.05). Reported hif-1 alpha is indicated as normalization relative median of OVCAR8 WT, OVCA429 WT, and OVCAR3 WT cells without treatment (**b**) (3 independent experiments; Unpair t-test; One-way ANOVA and Tukey’s multiple comparison test, p<0.05). (**c**) and (**d**) are confocal images showing hif-1 alpha during claudin-4 overexpression in OVCAR8 cells treated or not as indicated above at 96h. (**e**) and (**f**) are confocal images showing hif-1 alpha during claudin-4 downregulation in OVCA429 cells treated or not as indicated above at 96h. (**g**) and (**h**) are confocal images showing hif-1 alpha during claudin-4 downregulation in OVCAR3 cells treated or not as indicated above at 96h. Graphs shown median with 95% confidence interval (CI). Scale bar 10µm.

Notably, claudin-4 overexpression correlated with significantly more ROS production (**Figure 7a, left**) but not hif-1 alpha (**Figure 7b, left**), implying that claudin-4 overexpressing cells may have elevated basal ROS levels, potentially diminishing therapy efficacy(85) due to a protective effect of perinuclear F-actin (**Figure 2d**).(86,87) Supporting this, during claudin-4 downregulation, the positive relationship between claudin-4 overexpression and increased ROS generation was lost in OVCA429 cells (**Figure 7a, middle**). Unlike OVCAR8 overexpressing claudin-4 (**Figure 7a, left**), the ROS response to treatment was significantly higher in OVCA429 claudin-4 KD cells (**Figure 7a, middle**), which also exhibited reduced levels of perinuclear F-actin (**Figure 2e**). In contrast, claudin-4 downregulation in OVCAR3 cells led to increased ROS production (**Figure 7a, right**), with no changes in hif-1 alpha expression (**Figure 7b, right**) and stable perinuclear F-actin (**Figure 2f**). Although hif-1 alpha remained largely unaffected by claudin-4 overexpression or downregulation (**Figure 7b**), a significant increase was observed in hif-1 alpha levels following treatment in OVCA429 WT cells. This suggests potential differences in hif-1 alpha-mediated oxidative stress response between OVCA429 WT and OVCAR3 WT cells (**Figure 7b, middle and right**). Variations in ROS production between OVCA429 and OVCAR3 cells may be due to KRAS amplification in OVCAR3 cells, (19,54) which can enhance ROS generation in tumor cells.(88) Thus, the effect of our tripartite combination on inducing ROS could be masked by KRAS amplification.

Since claudin-4 modulation did not show significant changes in hif-1 alpha expression (**Figure 7b, at 4h**), we investigated its intracellular distribution using immunofluorescence. We observed notable changes in hif-1 alpha localization linked to claudin-4 modulation and extended treatment duration. Specifically, claudin-4 overexpression led to increased nuclear accumulation of hif-1 alpha, (**Figure 7c, d**) and higher protein levels at later time points (*See Supplementary Figure 6d*). Treatment further altered hif-1 alpha distribution and upregulated its expression in OVCAR8 cells (**Figure 7c, d**; *See Supplementary Figure 6d*). Claudin-4 downregulation in OVCA429 led to reduced hif-1 alpha levels (**Figure 7e, f** and *Supplementary Figure 6e*), while in OVCAR3 cells, it resulted in increased hif-1 alpha levels (**Figure 7g, h** and *Supplementary Figure 6f*). Notably, claudin-4 downregulation in OVCA429 cells caused hif-1 alpha to accumulate at the plasma membrane, an effect that was enhanced by treatment (**Figure 7e, f**) and partially mirrored in OVCAR3 cells (**Figure 7h**). These findings underscore a link between claudin-4 and hif-1 alpha, indicating that oxygen regulation and related factors, such as ROS,(60) play a role in claudin-4-mediated resistance to olaparib. They also suggest that targeting the oxidative stress response could enhance olaparib efficacy by disrupting claudin-4’s functional effects in ovarian cancer cells.

## DISCUSSION

In this study, we found that claudin-4 protects ovarian cancer cells by remodeling nuclear architecture and slowing cell cycle progression. This mechanism enables cancer cells to resist both the development of genomic instability and the effects of genomic instability-inducing agents like the PARP inhibitor olaparib. Therefore, targeting claudin-4 could reduce the dosage of olaparib required to induce cell death in ovarian cancer cells, thereby decreasing therapy resistance, possibly increasing patient survival.

Claudin-4 played a crucial role in regulating the dynamics of both nuclear lamina and the actin-cytoskeleton (**Figure 2**). Notably, claudin-4’s impact on nuclear architecture and the cytoskeleton was associated with nuclear constriction, suggesting that claudin-4 overexpression generates mechanical forces that shape the nucleus and prevent its enlargement (**Figure 4**). (3,44,89) This nuclear constriction may explain why cells overexpressing claudin-4 are more likely to be arrested in the G0-G1 phase and have more control over proceeding to the S-phase (**Figure 1a**). This suggests that claudin-4 may act as a brake, delaying the entry of ovarian cancer cells into the S-phase (**Figure 1d**). In contrast, cells with claudin-4 downregulation are more likely to be found in the G2-M phase (**Figure 1b**). Supporting this concept, lamin B1 has been reported to regulate the entry into the S-phase,(90) and our data indicated that claudin-4 modulation affected the nuclear localization of this protein. Specifically, claudin-4 caused the displacement of lamin B1 from the nucleus and promoted the stabilization of F-actin at the perinuclear and cell-to-cell regions (**Figure 3**). This phenotype could be linked to a reported exclusion mechanism mediated by fascin and actinin, which compete to bundle F-actin in different actin networks.(41,42) Overall, these results highlight claudin-4’s role in modulating the interplay between nuclear physiology and cell cycle progression, which may help reduce genomic instability formation. For example, claudin-4-induced reductions in genomic instability (**Figure 4**) contribute to therapy resistance, (23) which correlates with poor patient survival (*See Supplementary Figure 4a*). To investigate this mechanism further, we used the PARP inhibitor olaparib, an agent known to induce genomic instability as a mechanism to promote tumor cell death.(31,32) We observed that downregulation of claudin-4 was associated with a significant reduction in the concentration of olaparib required to inhibit the growth of OVCAR3 cells (**Figure 5c**), but not OVCA429 cells (**Figure 5b**). These differences underscore the heterogeneity among ovarian cancer cells and emphasize the importance of claudin-4 in modulating the response to PARP inhibitors in high-grade serous ovarian carcinoma (HGSOC), the most prevalent subtype of ovarian cancer, which accounts for over 75% of all ovarian cancer cases.(19,23,54) Importantly, targeting claudin-4’s functional effects in ovarian cancer cells using CMP and FSK—potentially involving alterations in LAT1 and lamin B1 (**Figure 6a-c**) (*See Supplementary Figure 5d-i*)—in combination with olaparib, led to greater reductions in all ovarian cancer cell survival tested at the lowest concentrations of olaparib (**Figure 5j-l**).

As ovarian cancer cells were cultured for 7 days in the colony formation assay, we speculated that hypoxia may play a role in claudin-4-mediated resistance to olaparib. This hypothesis is supported by the close link between claudin-4 and hif-1 alpha in regulating hypoxia through a feedback mechanism that may affect autophagy,(91) and a similar association between LAT1 and hif-1 alpha under these conditions.(92,93). Our evaluation of claudin-4’s role in therapy resistance revealed that its effect was associated with previously reported biological interactions, including LAT1(19) (**Figure 5d-f**), hif-1 alpha(91) (**Figure 7b-h**), (91) and lamin B1 (**Figure 5d-f, bottom**), in potential cellular processes such as autophagy, hypoxia, and nuclear lamina remodeling, respectively. This is particularly evidenced with evaluation of an oxidative stress response (**Figure 7**) during the triple combination treatment. This treatment resulted in significant increase of ROS and hif-1 alpha in all ovarian cancer cells (**Figure 7**), suggesting that ovarian cancer cells generate an oxidative stress response to counteract olaparib treatment,(60,81) potentially through changes in the metabolism of mitochondria.(94) Since an excessive oxidative stress response can lead to cell death(95) and the nuclear lamina can protect against ROS,(84) these data suggest that the claudin-4’s role in remodeling the nuclear architecture (**Figure 2**) may help protect tumor cells during excessive oxidative stress response during olaparib treatment. Consequently, modulating the oxidative stress response could further potentiate the combined olaparib/FSK/CMP in reducing ovarian cancer cell survival. For instance, previous studies have shown that FSK and metformin can decrease oxidative stress, (96) which could interfere with the response of ovarian tumor cells to the tripartite treatment (**Figure 7**). Additionally, the combination of metformin with olaparib inhibits proliferation of ovarian cancer cells.(97) Our results highlight the potential of this tripartite treatment strategy to reduce therapy resistance *in vivo*.

## ACKNOWLEDGEMENTS

We acknowledge philanthropic contributions from D. Thomas and Kay L. Dunton Endowed Chair in Ovarian Cancer Research, the McClintock-Addlesperger Family, Karen M. Jennison, Don and Arlene Mohler Johnson Family, Michael Intagliata, Duane and Denise Suess, Mary Normandin, and Donald Engelstad. In addition, we acknowledge the Cancer Center Support Grant (P30CA046934) and our Flow Cytometry Shared Resource, University of Colorado Anschutz | Medical Campus.

## Author’s contribution

BB and MN conceived the overall project. BB supervised the research project. FRV performed the experiments and data analysis along with all authors. FRV wrote the manuscript with the contribution of all authors.

## Abbreviations

CMP: claudin mimic peptide
HGSOC: high-grade serous ovarian carcinoma
LGSOC: low-grade serous ovarian carcinoma
MOC: mucinous ovarian carcinoma
PARP: Poly (ADP ribose) polymerase
FSK: forskolin
ROS: reactive oxygen species.

## SUPPLEMENTARY FIGURES

**Supplementary Figure 1.**
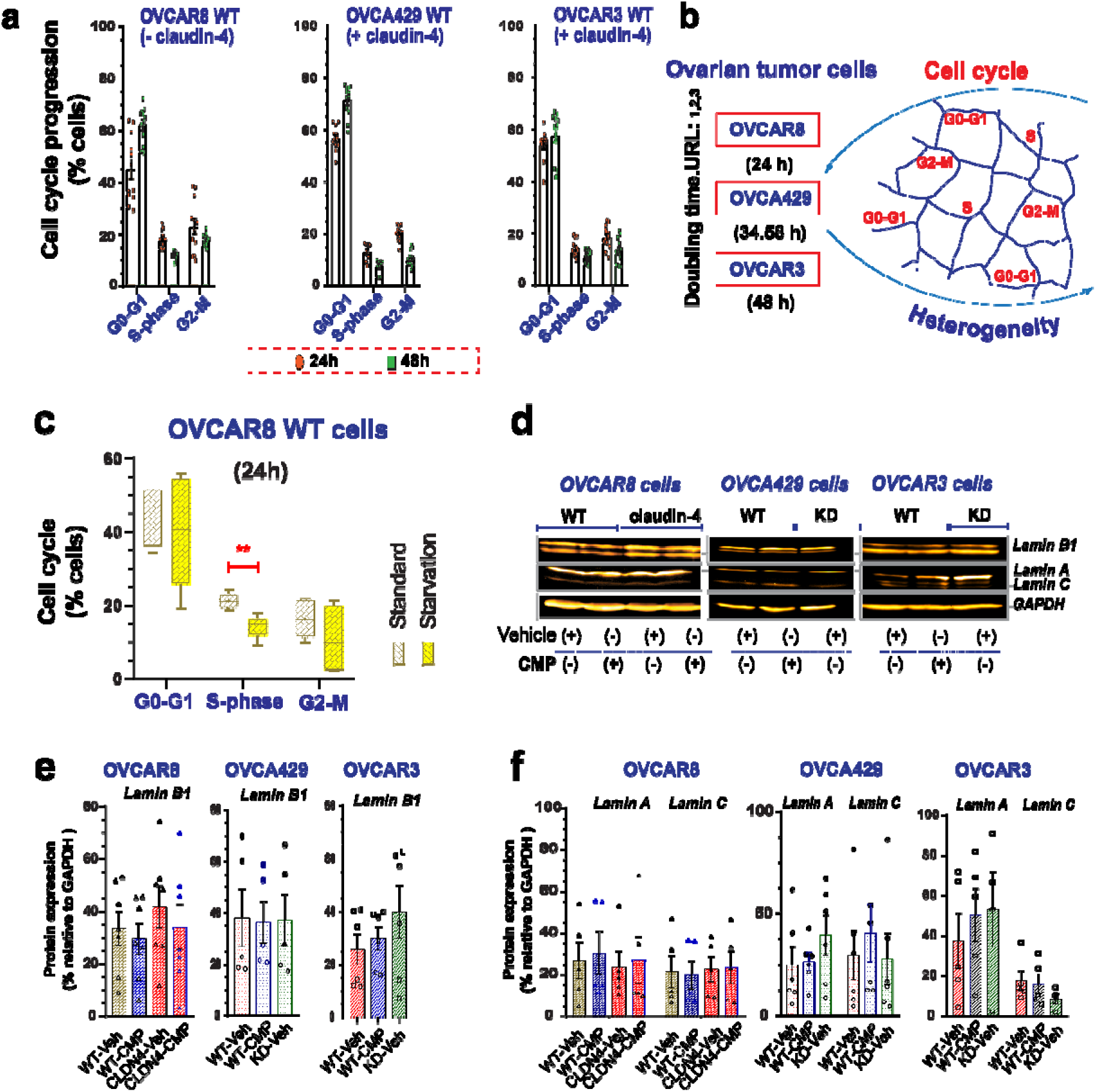
**(a)** Graphs show WT ovarian tumor cells in different phases of the cell cycle (green symbol at 24h; red symbol at 48h) which change over time. (**b**) Drawing highlighting heterogeneity of ovarian tumor cells, indicated as localization in different phases of cell cycle (references: URL 1-3). Also, it highlights differences in doubling time among different ovarian tumor cells. (**c**) Quantification of cell cycle progression by flow cytometry of propidium iodide-stained OVCAR8 WT cells during standard (RPMI, 10% FBS) culture conditions or starvation (RPMI, 1% FBS). (**d**) immunoblotting for lamin B1 and lamin A/C and corresponding quantification (**e**, lamin B1; **f**, lamin A/C) relative to loading control (4 independent experiments), respectively. (One-way ANOVA and Tukey’s multiple comparisons test; p<0.5). Graphs show mean and SEM, scale bar 5µm.

**Supplementary Figure 2.**
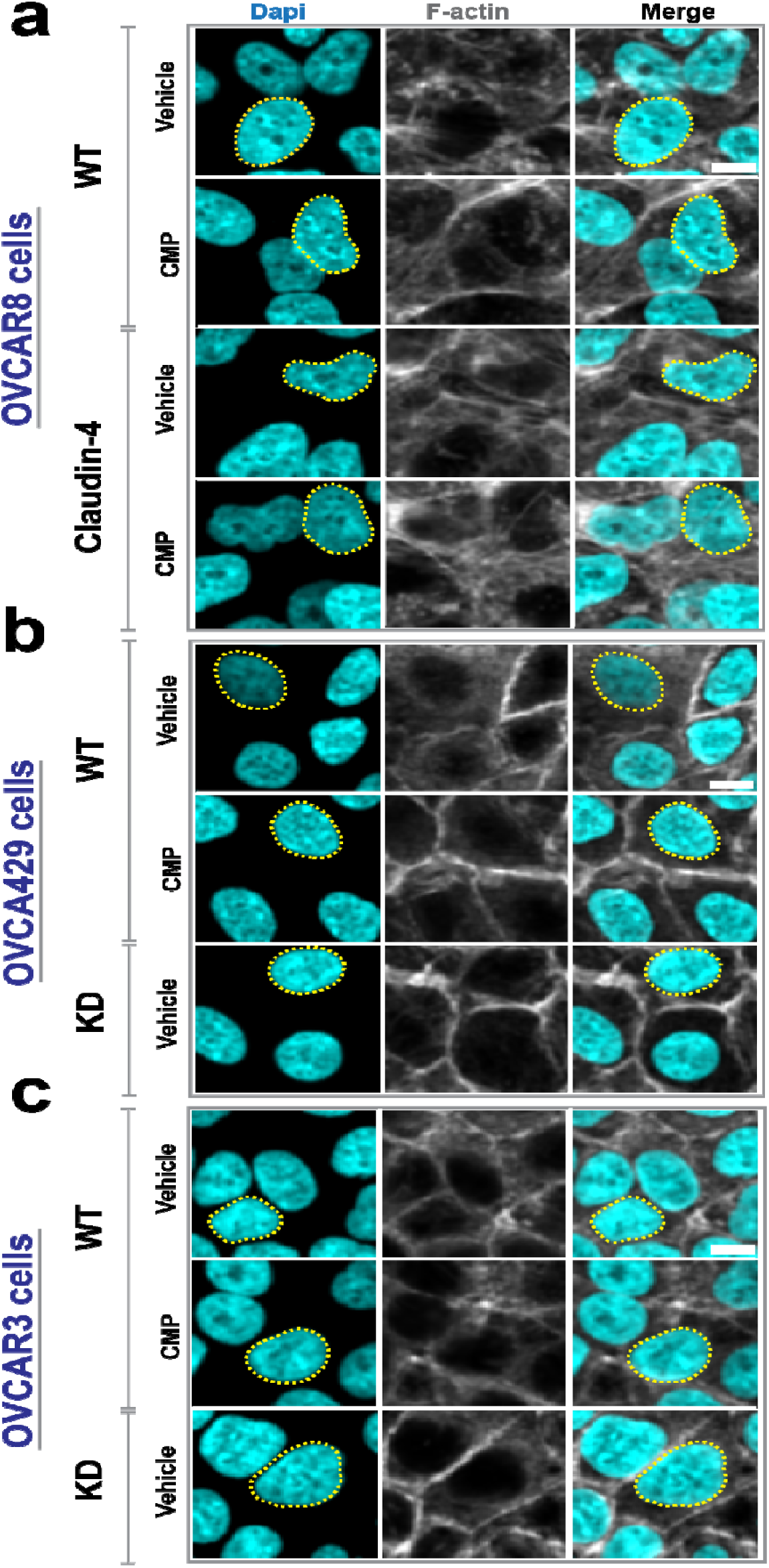
Epithelial ovarian cancer cells were treated with CMP (400µM) for 48h. Subsequently, cells were stained with dapi (nuclei) and phalloidin (F-actin) to carry out a morphometric characterization. (**a**), (**b**), and (**c**) show representative confocal images of OVCAR8, OVCA429, and OVCAR3 cells, respectively. Scale bar 10µm.

**Supplementary Figure 3.**
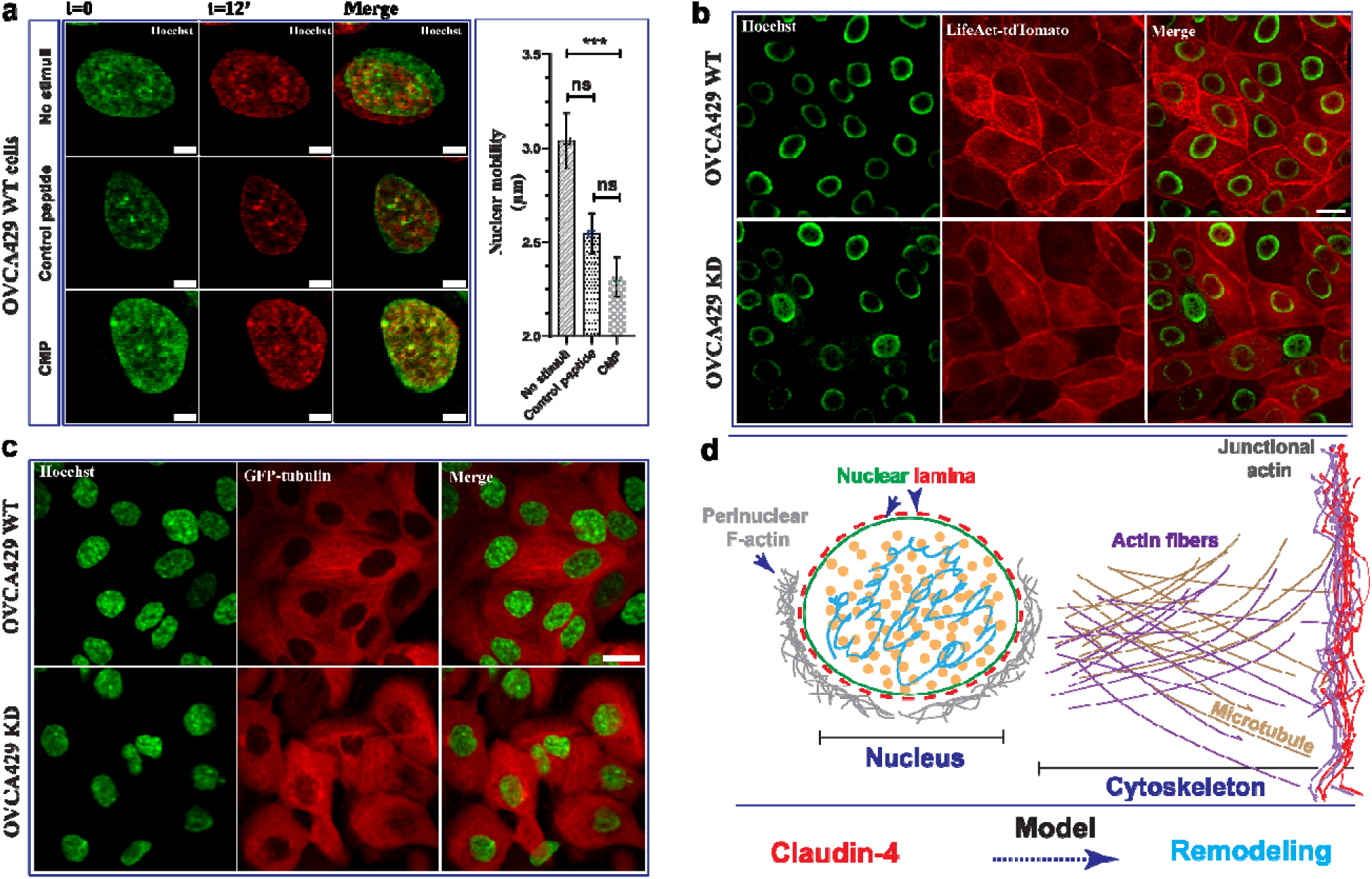
Live cell imaging evaluation of ovarian tumor cells. (**a**) shows selected confocal images (xyt/30min/37°C) of nucleus at different time points (cells treated with CMP or a control peptide; 400µM for 24h), which were overlaid to highlight the nuclear mobility; right, quantification of nuclear mobility (n= no stimuli, 44 cells; control peptide, 44 cells; CMP, 41 cells; 4 independent experiments; Kruskal-Wallis test and Dunn’s multiple comparison test, p<0.5). (**b**) shows representative confocal images of living cells expressing LifeAct-tdTomato (marker of F-actin) without any stimuli. (**c**) shows representative confocal images of living cells (xyt; maximum projections) of cells expressing GFP-tubulin without any stimuli. (**d**) presents a model that highlights our findings on the role of claudin-4 in the remodeling of the nucleus and cytoskeleton. Nuclear remodeling was linked to alterations in lamin B1 localization, resulting in changes to the nuclear lamina, along with the accumulation of perinuclear F-actin. Cytoskeletal remodeling involved changes in actin fibers, the microtubule network, and junctional F-actin, which supports cell-to-cell junctions, potentially impacting nucleus positioning, cell cycle, and cell morphology. Graph shows mean and SEM, scale bar 20µm.

**Supplementary Figure 4.**
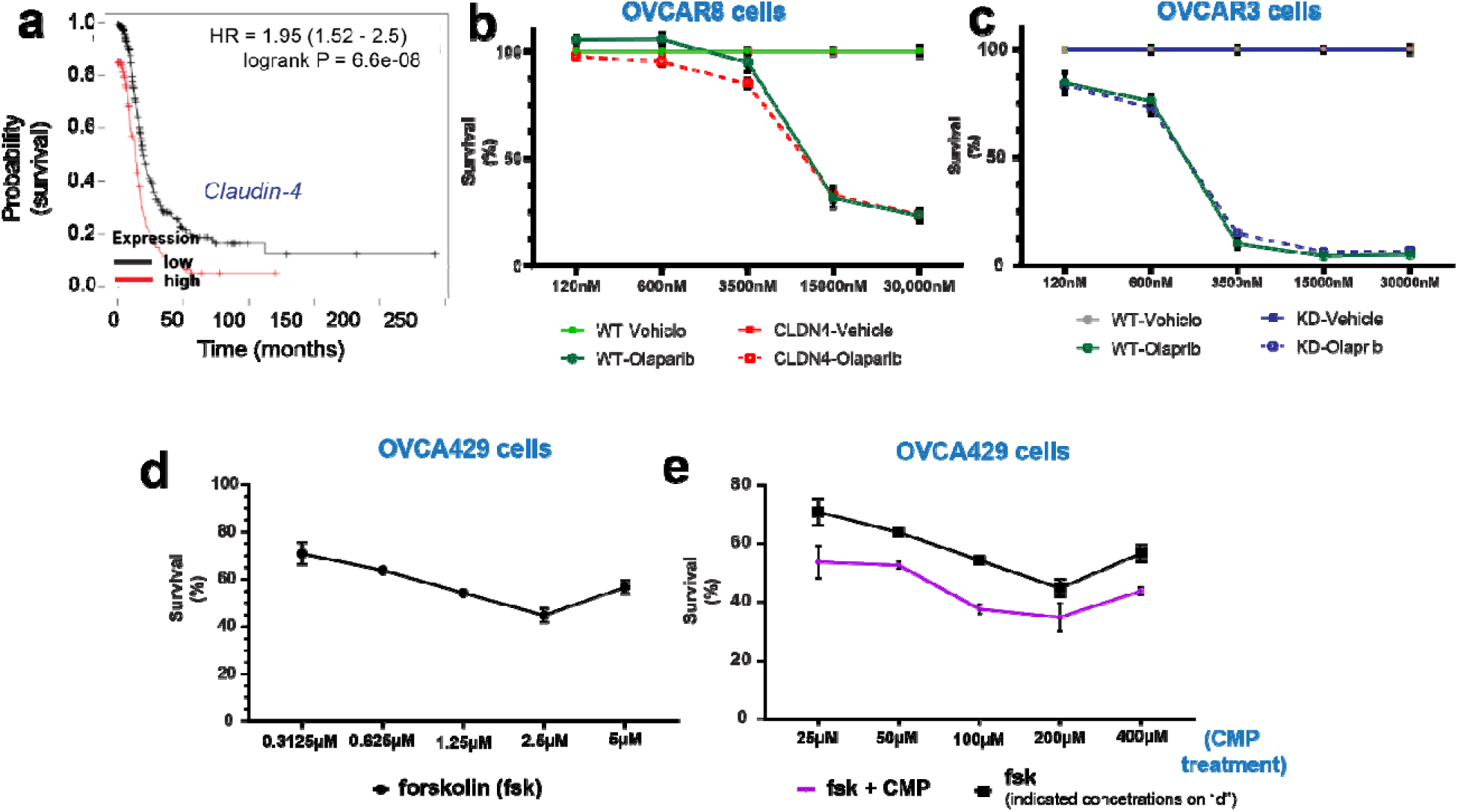
(**a**) shows a Kaplan-Meier curve based on claudin-4 expression in human ovarian tumors (Kaplan-Meier Plotter) highlighting the association of higher claudin-4 expression with reduced patient survival. (**b**) and (**c**) show survival of ovarian tumor cells treated with olaparib during overexpression (OVCAR8 cells) and downregulation (OVCAR3 cells). (**d**) and (**e**), show percentages of ovarian tumor cell survival during various concentrations of fsk and a comparison with a combination of fsk at 5µM with various concentrations of CMP, respectively. Graphs show mean and SEM.

**Supplementary Figure 5.**
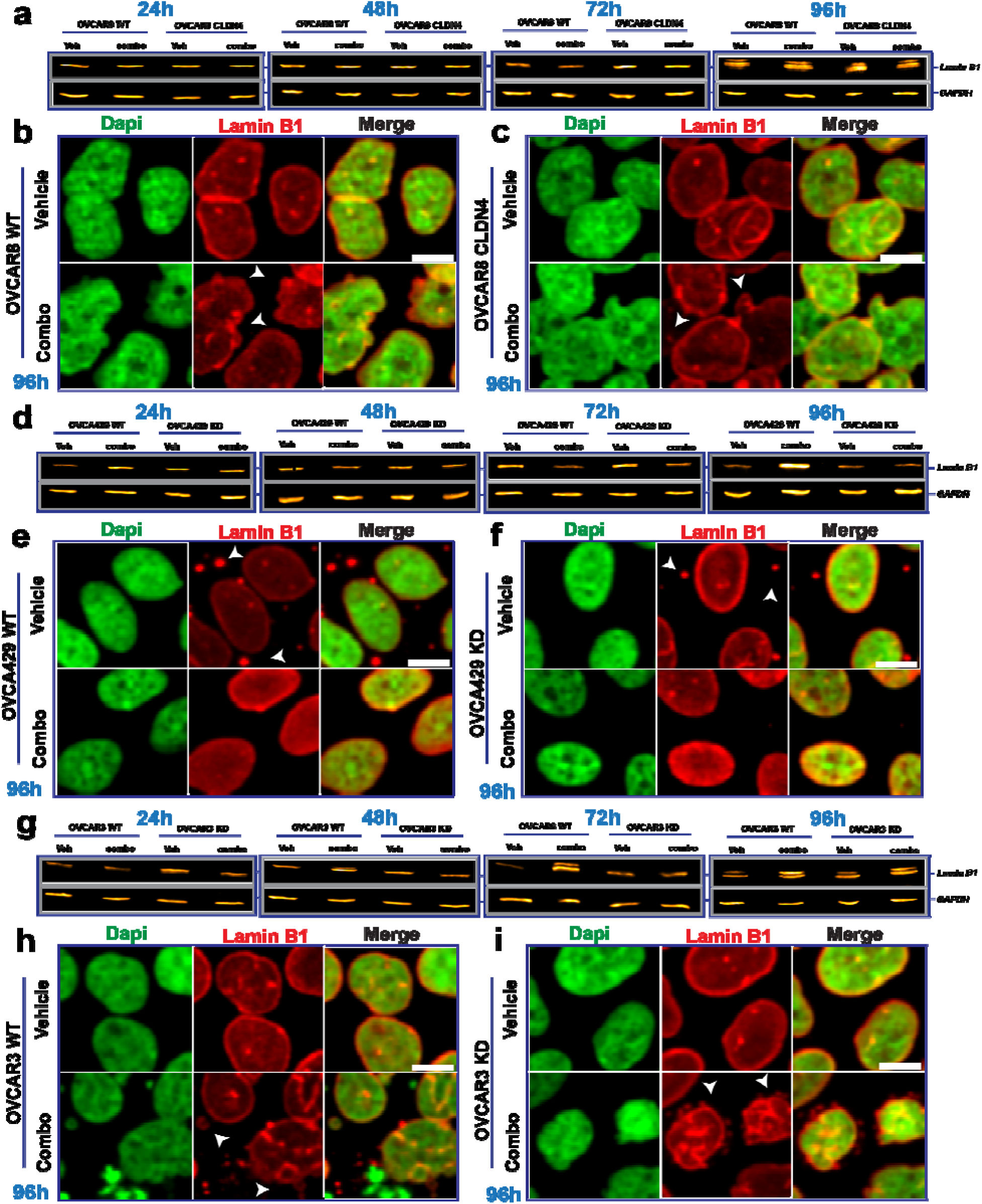
(**a**), (**b**), and (**c**) show lamin B1 protein expression during a tripartite combination treatment of olaparib (600nM), fsk (5µM), and CMP (400µM) at different time points in OVCAR8, OVCA429, and OVCAR3 cells, respectively. (**d**) to (**i**) are confocal image of ovarian tumor cells showing the intracellular distribution of lamin B1 during the same treatment at 96h for OVCAR8, OVCA429, and OVCAR3, respectively. Scale bar 10µm.

**Supplementary Figure 6.**
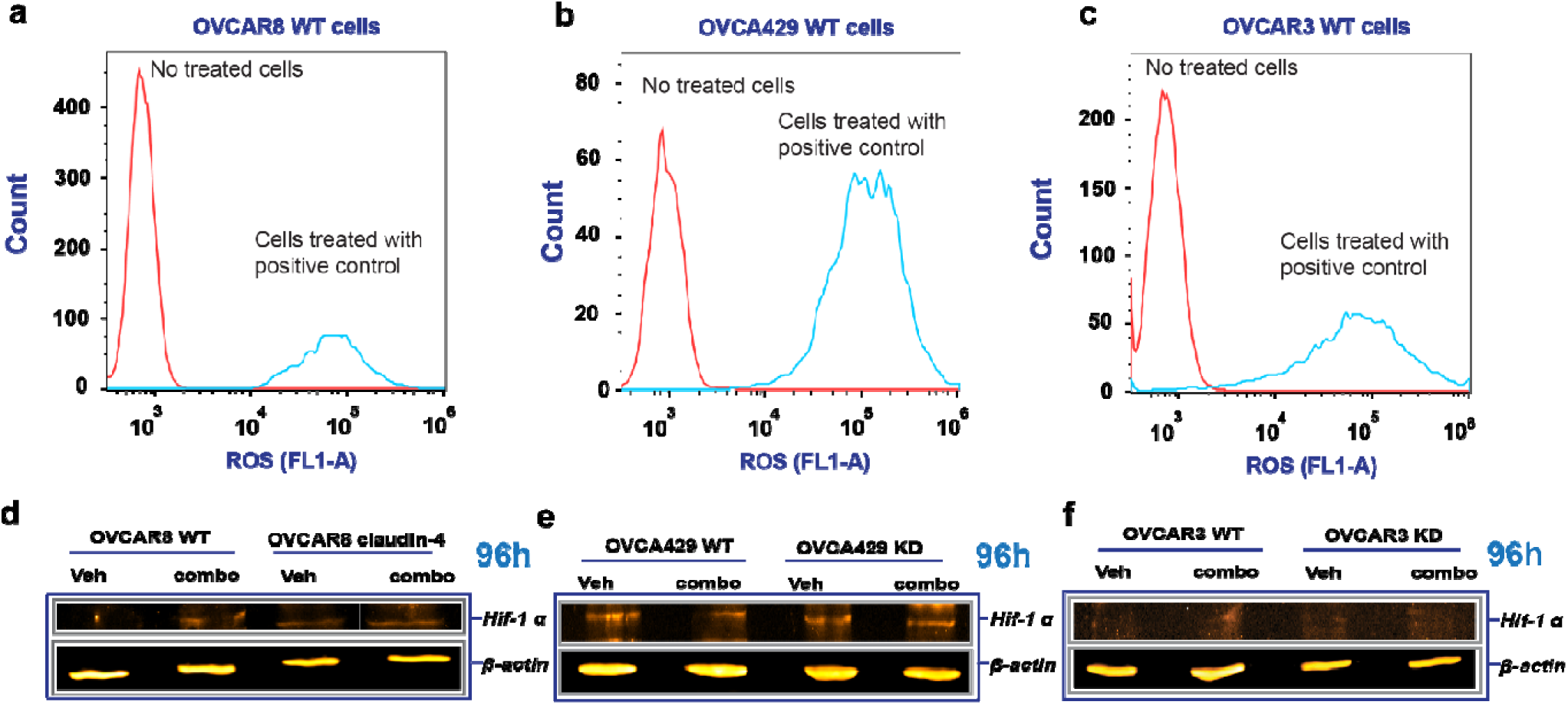
(**a**), (**b**), and (**c**) are histograms confirming reactive oxygen species (ROS) generation by OVCAR8, OVCA429, and OVCAR3 cells, respectively, using TBHP at 250µM as a positive control. (**d**), (**e**), and (**f**) show hif-1 alpha protein expression during a tripartite combination treatment of olaparib (600nM), fsk (5µM), and CMP (400µM) at 96h of treatment in OVCAR8, OVCA429, and OVCAR3 cells, respectively.

